# Hippocampal theta sequences in REM sleep during spatial learning

**DOI:** 10.1101/2021.04.15.439854

**Authors:** Mark C. Zielinski, Justin D. Shin, Shantanu P. Jadhav

## Abstract

Rapid eye movement (REM) sleep is known to play a role in hippocampally-dependent memory, yet the activity and development of hippocampal neuronal ensembles during this state is not well understood. Here we investigated patterning of CA1 place cell activity by theta oscillations, a shared electrophysiological hallmark of both waking behavior and REM sleep, in male rats learning a spatial memory task. We report the existence of REM theta sequences, sequential reactivations of place cells in REM theta that parallel waking theta sequences. REM and wake theta sequences develop rapidly with experience, recapitulating behavioral sequences of compressed space in forward and reverse directions throughout learning. REM sleep exhibited a balance of forward and reverse sequences in contrast to predominantly forward wake theta sequences. Finally, we found that a CA1 neuronal population known to shift preferred theta phases in REM exhibited differential participation in wake and REM theta sequences. In particular, this phase-shifting population showed an increased contribution to REM theta sequence representations after behavioral performance asymptotes and the task is learned, supporting a previously hypothesized role in depotentiation. These findings suggest a role for REM associated theta sequences in state dependent memory functions of the hippocampal circuit, providing evidence that REM sleep is associated with sequence reactivation that can support consolidation of representations necessary for memory guided behavior.

## INTRODUCTION

Episodic memory guides behavior in the face of a continuously changing environment, integrating new information into existing representations of the external world or erasing old memories and creating new ones. The hippocampus is necessary for encoding and consolidation of memories from their labile form into stable long-term representations (Müller and Pilzecker, 1900; Dudai, 2004; Squire et al., 2015), with existing memory traces either updated, generalized, or forgotten (Alberini, 2005; Li et al., 2017; Izawa et al., 2019). Hippocampal place cells and their inputs support spatial and contextual memories (O’Keefe, 1976; Eichenbaum and Cohen, 2004; Dupret et al., 2010), by individually representing components of specific episodes (Moser et al., 2008) or sequences (Eichenbaum et al., 1999; Pastalkova et al., 2008; Tingley et al., 2018) in space and time. Sequential activation of place cells at fast timescales within single theta cycles and sharp-wave ripples can recapitulate this behavioral neural activity in a temporally compressed manner during both waking and offline states (Skaggs and McNaughton, 1996; Lee and Wilson, 2002; Karlsson and Frank, 2009; Jadhav et al., 2012; Drieu and Zugaro, 2019). These sequences are ideal for binding representations at a cellular timescale fast enough to trigger long term potentiation or depression (Mizumori et al., 1990; Bi and Poo, 1998), with waking and rest states thought to separate encoding and retrieval from consolidation and forgetting, respectively (Buzsáki, 1989; Mizuseki and Miyawaki, 2017; Poe, 2017; Samanta et al., 2020).

Coordination of hippocampal ensembles (Winson, 1978) during exploration, movement, and active sensation is patterned by the theta rhythm, a regular 6-12 Hz rhythm driven by the medial septum (Petsche et al., 1962) and seen in hippocampally associated areas during waking (Vanderwolf, 1969) and REM sleep (Jouvet, 1969; Buzsáki, 2002; Montgomery et al., 2008; Grosmark et al., 2012). The phase of spikes and ensemble firing relative to theta can directly influence neuronal plasticity across these areas (Huerta and Lisman, 1993, 1995; Hölscher et al., 1997; Orr et al., 2001; Griffin et al., 2004), and also induce bidirectional changes in memory and behavior (Pavlides et al., 1988; Hyman et al., 2003; Siegle and Wilson, 2014). The most well-known influence of this rhythm on individual place cells is the phenomenon of phase precession, where spikes shift their preferred firing relative to the phase of theta over the extent of a place field in an experience dependent manner (O’Keefe and Recce, 1993; Skaggs et al., 1996; Harris et al., 2002; Schmidt et al., 2009). Theta driven sequences emerge at the perspective of multiple place cells (Skaggs et al., 1996; Dragoi and Buzsáki, 2006; Foster and Wilson, 2007), are experience-dependent and separable from precession (Foster and Wilson, 2007; Feng et al., 2015; Silva et al., 2015), have been shown to map behavior in the forward and reverse directions (Cei et al., 2014; Wang et al., 2020), extend dynamically to goals (Gupta et al., 2012; Wikenheiser and Redish, 2015b), and serially represent options in deliberative decision making (Johnson and Redish, 2007; Gupta et al., 2012; Papale et al., 2016; Redish, 2016; Kay et al., 2020). Although theta oscillations are prominent in REM sleep, whether this theta in REM sleep organizes place cells into sequences in a similar manner as its waking counterpart is currently an open question.

REM sleep is associated with a unique neuromodulatory landscape and has been proposed to have roles in memory processes, specifically both consolidation and forgetting (Rasch and Born, 2013; Poe, 2017; Samanta et al., 2020). The theta oscillation itself undergoes a change in REM sleep, reversing its direction of flow through the hippocampal formation, from the subiculum to CA1 to CA3 (Jackson et al., 2014; Genzel et al., 2015). REM also induces transient changes in synapse maintenance and formation, cellular excitability, and ensemble synchrony (Montgomery et al., 2008; Grosmark et al., 2012; Li et al., 2017). Superficial and deep CA1 neurons are modulated by REM differently, with deep CA1 cells shifting their preferred theta phases from wake to REM sleep (Poe et al., 2000; Mizuseki et al., 2011; Valero et al., 2015; Wang et al., 2020). Reverberations of waking activity in REM sleep, at the same timescale as behavioral activity, have also been reported (Louie and Wilson, 2001; Peyrache et al., 2015), though the development and detailed properties of this reactivation is unknown. Experience and novelty dependent changes in REM theta are thought to occur from these and other lines of evidence, but recent findings have been mixed (Poe et al., 2000; Mizuseki et al., 2011). Further, despite the similar network states in REM and wake, and the role that theta plays in episodic memory, little is still known about the existence of compressed hippocampal theta sequences in REM, their functional properties and development, and their potential role in behavior. We therefore investigated this question using continuous hippocampal CA1 ensemble recordings across interleaved behavior and rest sessions in rats learning a spatial alteration task in a single day.

## RESULTS

### Trajectory representation and REM sleep during single-day acquisition of a W-track spatial alternation behavior

In order to address the potential role that theta associated place cell sequences have in REM sleep, we monitored hippocampal ensembles *in vivo* throughout single-day acquisition of a W-track spatial alternation task known to require working memory (**Figure 1A,B**) (Jadhav et al., 2012; Maharjan et al., 2018; Shin et al., 2019). The W-track continuous alternation task requires animals to visit reward ports situated at the end of the three track arms in a specific behavioral sequence. Correct inbound or outbound trials are rewarded in the following manner: starting from the center arm, the animal must visit an outer arm reward well (outbound trajectory), then return from the outer arm well to the center arm well (inbound trajectory) to consume reward. All subsequent outbound trials require the animal to recall the previous outer arm visit, and visit the opposite outer arm well that was not previously chosen. This sequence then continues, with center arm visits interleaved with alternating outer arm visits (**Figure 1A, left**). Run sessions consist of 15-20 minutes of exposure to this track, followed by a ∼30 min rest session in a familiar sleep box (**Figure 1A, right**).

**Figure 1.**
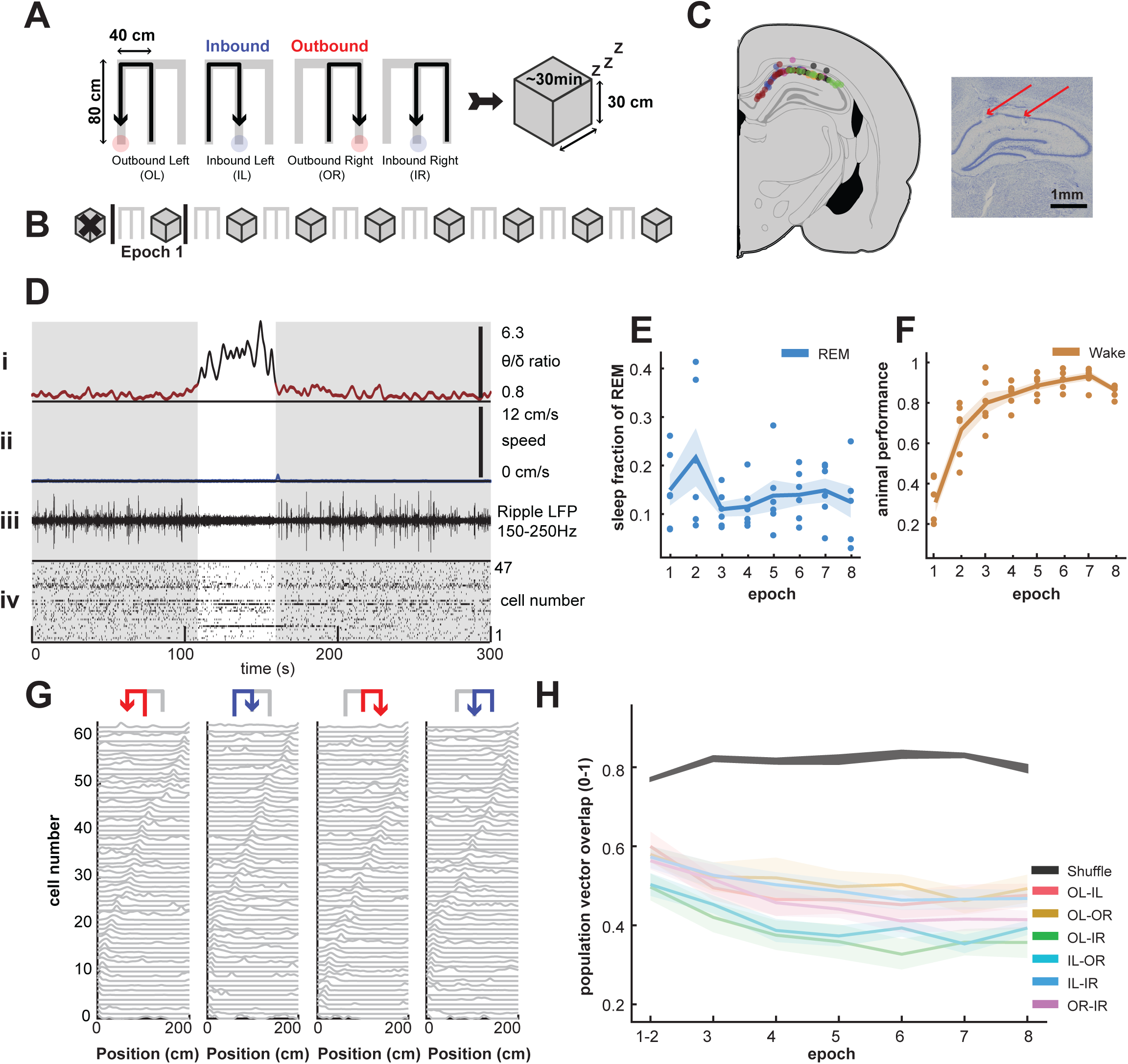
REM sleep and distinct behavioral trajectory representations in CA1. **(A)** Schematic of one behavioral epoch; a W track session (Run/Wake) and subsequent sleep session (Sleep/REM). A successful behavioral sequence in the W track spatial alternation task requires alternation between outbound arm visits (red wells) and returns to the center arm (blue wells) for a total of four trajectories depicted from left to right: Outbound Left (OL), Inbound Left (IL), Outbound Right (OR), and Inbound Right (IR). **(B)** Illustration of the single day W track acquisition task, consisting of eight behavioral epochs (one run session, followed by one sleep session) over the course of one day. An initial sleep session prior to the first track exposure was not used. **(C) (Left)** Schematic of all hippocampal tetrode locations in all animals, one color per animal (n=6 animals). **(Right)** Coronal brain section showing locations of two CA1 tetrode lesions (red arrows). **(D)** Segmentation of SWS and REM during sleep sessions. **(i)** shows the theta delta ratio for a given sleep session, with theta/delta ratio greater than 2 highlighted in black. Animal speed in **(ii)** is consistently immobile or below a threshold of 3 cm/s for all REM sessions. Ripple power is consistently low in REM sleep **(iii)**, with no ripple events exceeding 3sd. Note the change in overall spike rate and place cell activity in **(iv)**. **(E)** Sleep fraction of REM (blue) does not change over behavioral epochs (all p> 0.81, Kruskal-Wallis test with Tukey-Kramer correction for multiple comparisons). **(F)** Animal performance in wake sessions (orange), calculated as the total number of correct inbound and outbound trials over the total number of trials. Animal performance asymptotes ∼ behavioral epoch #5, defined as the first behavioral epoch where performance is significantly different from the first behavioral epoch (p< 0.01 for behavioral epoch comparison 1 vs. 5, Kruskal-Wallis test with Tukey-Kramer correction for multiple comparisons). **(G)** Firing rate curves of all place cells for one animal in one behavioral epoch along a linearized trajectory (0-200cm, normalized to peak rates). From left to right, cells are sorted according to peak place field locations on that trajectory. **(H)** Population vector overlap (PVO) correlations of place cells between each trajectory for every behavioral epoch (light), and the corresponding 95% CIs of trial permuted shuffles (dark regions shows shuffles for all trajectory comparisons). All trajectory templates are distinct from one another throughout the experiment (all p < 0.01, permutation tests). Epochs 1 and 2 were combined due to low number of trials in the first epoch during permutation. Outbound Left (OL), Inbound Left (IL), Outbound Right (OR), and Inbound Right (IR). Data are means (solid lines) ± SEM (shaded area). \**P* < 0.05; ns, not significant.

Custom adjustable microdrive arrays were used to target tetrodes to dorsal CA1 regions (see **Methods**, **Figure 1C**). Electrophysiological recordings monitored activity of hippocampal ensembles throughout the entirety of the experiment over task acquisition (5.5-6.5 hours). A behavioral epoch consisted of one run session, followed by one sleep session, for a total of 8 behavioral epochs (**Figure 1B**, an initial sleep session prior to track exposure was not used for this analysis). Place cell templates in a given run session were used for subsequent decoding in the following sleep session. All animals exhibited rapid learning of the task (**Figure 1F**), and the proportion of correct overall trials, and therefore learning, was asymptotic by behavioral epoch 5 (Kruskal–Wallis with Tukey-Kramer correction for multiple comparisons between the 8 behavioral epochs; X^2^(7)=31.81, comparison between epochs 1 and 5 significantly different, p = 0.0096; all comparisons between epochs 5-8 were not significantly different, p > 0.76). This experimental design thus allowed for investigation of CA1 dynamics in waking and sleep states over the course of learning, from initial acquisition to later correct performance (mean ± SEM number of hippocampal place cells per animal epoch; wake, 47 ± 1.5 cells; sleep, 40 ± 1.2 cells).

REM sleep in rodents is fragmented, with changes in LFP rhythms during sleep reminiscent of wake, namely increase in theta (6-12 Hz) power and decrease in delta (1-4 Hz) power (Twyver, 1969; Rasch and Born, 2013). Using established methods for REM detection ((Grosmark et al., 2012; Kay et al., 2016; Tang et al., 2017), see **Methods** and **Figure 1D**), namely immobility, lack of ripple power corresponding to sharp wave ripple events, and high theta to delta ratio, the fraction of REM sleep during rest sessions was consistent across all behavioral sessions (**Figure 1E**, Kruskal–Wallis with Tukey-Kramer correction for multiple comparisons between the 8 behavioral epochs; X^2^(7)=3.69, p = 0.8152, all comparisons p > 0.8050), in agreement with prior literature (Twyver, 1969; Mendelson and Bergmann, 1999).

CA1 place cells are direction and trajectory selective, with distinct spatial encoding of all 4 trajectories (outbound left, outbound right, inbound left, and inbound right) in the W-track task (Shin et al., 2019). **Figure 1G** shows normalized firing rates of all place cells for a single animal and epoch, sorted according to place field peak in the respective trajectory. These ensembles are directionally selective from the onset of behavior and direction selectivity develops rapidly with experience, as previously reported (Frank et al., 2004; Foster and Wilson, 2006; Shin et al., 2019; Xu et al., 2019). Further, as we have previously shown (Shin et al., 2019), trajectory specific sequences of place cell ensembles were unique and differentiable from one another for each animal and behavioral epoch (**Figure 1H**, permutation test, all comparisons between real and shuffled data; p<0.01), allowing for unambiguous detection, identification, and tracking of potential forward and reverse sequences throughout behavior. The stability of REM sleep fraction during single day learning, in conjunction with monitoring of hippocampal ensembles, enabled us to examine the development of hippocampal ensemble activity patterns in waking and REM states.

### Theta properties and parameters in wake and REM sleep

We first examined theta-phase associated properties of place cells in the two states, wake and REM Sleep. In particular, since reversal of preferred theta phase in REM sleep is known to occur for a subset for place cells (Poe et al., 2000; Mizuseki et al., 2011), we examined the prevalence of REM theta phase reversal during spatial alternation learning. We found large fractions of neurons significantly phase locked to theta (Rayleigh z test for non-uniformity p < 0.05) in wake and REM sessions (mean ± SEM phase locked cells per animal epoch; Wake 61% ± 8.9% (30 ± 4 cells); REM sleep 37% ± 5.3% (15 ± 2 cells)). The distributions of mean preferred phases in wake and REM had a small but significant difference (**Figure 2A**, Kuiper two-sample test; the two distributions for preferred phases of the phase-locked neurons are significantly different, T(Kuiper) = 1.3e04, p = 0.005), in agreement with previous literature (Poe et al., 2000; Mizuseki et al., 2011). To investigate whether a subset of hippocampal neurons in this task also reversed their preferred phases, we analyzed place cells which had significant phase-locking in both wake and REM sleep (per animal mean ± SEM; 30 ± 5 cells). We found that a fraction of place cells shifted their preferred phases in REM sleep (**Figure 2B**; over all animals and behavioral epochs, 30% ± 2.3% (9 ± 1 cells), mean ± SEM), using criteria (see **Methods**) similar to previously established guidelines delineating shifting cells from non-shifting cells (Mizuseki et al., 2011). Individual examples of two cells in **Figure 2C** show these phase preference shifts between Wake and REM.

**Figure 2.**
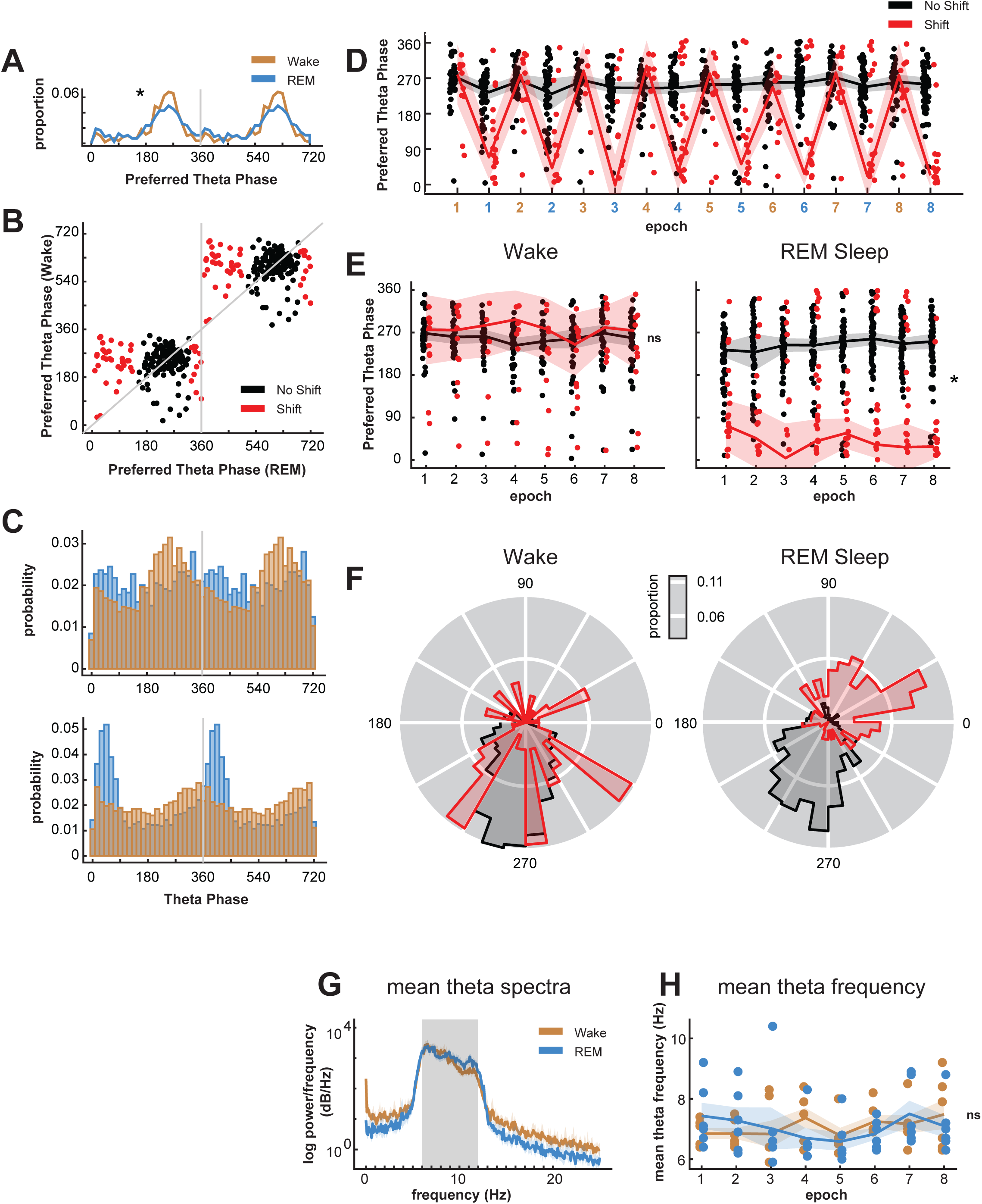
Theta phase preference in wake and REM. **(A)** Distribution of mean preferred theta phases of all place cells that showed significant theta-phase modulation (p<0.05, Rayleigh’s z-test) in wake (orange) and REM sleep (blue). The two distributions of preferred phases differ in run and REM sleep (p < 0.005, Kuiper’s test). **(B)** Distribution of mean preferred theta phases for both run and REM sleep. Only neurons which had at least one epoch of significant theta phase modulation in both run and REM are shown. Neurons with <=140° or >=320° preferred phases centered around peak preferred phase during REM sleep are designated as shifting cells (red), while the remainder are considered non shifting cells (black), similar to (Mizuseki, 2011). **(C)** Example individual phase shifting cell examples, with histograms of spike theta phases in wake (orange) and REM sleep (blue). **(D)** Distribution of preferred theta phases of all place cells that showed significant theta-phase modulation per behavioral epoch, showing both shifting (red) and non-shifting (black) cells. Data are circular means (solid lines) ± bootstrapped 95% CIs (shaded area). Note that the circular Y-axis wraps around at 0 and 360. **(E)** Same as in **C**, split into wake **(left)** and REM sleep **(right)** epochs. **(Left)** Preferred phase of shifting and non-shifting cells during wake is similar for all epochs (p = 0.52, Watson-Willams test). **(Right)** Preferred phase of shifting and non-shifting cells is different over all epochs in REM (p < 1e-99, Watson-Willams test). Data are circular means (solid lines) ± bootstrapped 95% CIs (shaded area). \**P* < 0.05; ns, not significant. **(F)** Same as in **E**, summarized across all epochs as polar plots, split into wake **(left)** and REM sleep **(right)** sessions. **(G)** Mean theta spectra in wake and REM for all animals. **(H)** Mean theta frequencies for all animals do not differ in wake vs. REM sleep (p = 0.4080, Friedman Test). Data in **(G)-(H)** are means (solid lines) ± SEM (shaded area). \**P* < 0.05; ns, not significant.

In light of this notable fraction, we next examined theta phase-locking and phase-shifting throughout the duration of learning. **Figure 2D** shows the preferred theta phases over all epochs for shifting and non-shifting cells, with interleaved wake and REM sleep sessions. Mean preferred theta phases split by shifting and non-shifting cells show a clear transition from wake to REM sleep for the shifting cells for all epochs. In wake, preferred theta phases are statistically similar between shifting and non-shifting cells (Watson-Williams test, p = 0.52), in agreement with previous findings (Mizuseki et al., 2011) (**Figure 2E, left**). Preferred phases in REM are significantly different between the two populations (Watson Williams test, p < 1e-99) (**Figure 2E, right**). This shifting effect across all epochs is summarized as polar plots in **Figure 2F** for wake **(left)** and REM **(right)**. The number of phase-shifting cells did not change significantly across epochs during single day learning (there was no statistically significant difference in proportion of wake and REM phase shifting cells over epochs, χ2(7) = 12.69, p = 0.94; χ2 test of independence). We also examined properties of theta oscillations in the two states and found an overall similarity in theta frequency content (6-12 Hz, see **Methods**) across wake and REM in all animals (**Figure 2G**), with mean theta frequencies similar across the two behavioral states throughout the course of behavior (**Figure 2H**; there was no statistically significant difference between wake and REM theta frequencies, χ2(1) = 0.6847, p = 0.4080. Friedman Test).

### Detecting theta sequences in wake and REM sleep

Next, in order to assess the existence, content, and evolution of theta sequences, we used established methods to detect candidate periods of theta cycles during behavior and REM (see *High-theta segmentation and REM state identification,* and *Theta cycles and decoding* in **Methods** for details), and then used a Bayesian decoding approach to identify sequential reactivation events (Zhang et al., 1998; Davidson et al., 2009; Feng et al., 2015). Theta cycles, and therefore sequences, were segmented from trough to trough in waking behavior (Johnson and Redish; Gupta et al., 2012; Feng et al., 2015; Wu et al., 2017). Since there is a prevalence of phase reversing cells in REM sleep which preferentially fire during theta trough (**Figure 2**), as well as the known reversal of activity flow of the theta rhythm in REM (Jackson et al., 2014; Genzel et al., 2015), we segmented theta cycles from peak to peak in REM sleep. This also allowed us to assess the contribution of phase-reversing cells to potential theta sequences in REM. **Figure S1A** shows LFP, theta phase, spikes, animal position, and decoding for a 0.5 second period of behavior on the W-track task, with decoded track position for each of the four trajectories. **Figure S1B** shows the same for an equivalent segment of REM sleep, using place cell templates from the preceding run session.

Sequential structure of reconstructed position during reactivation is evaluated primarily by a trajectory score measure, which assesses a decoded Bayesian posterior for replay-like structure by finding the optimal virtual slope and intercept that maximizes an objective function - in this case, the total amount of decoded probability captured within the boundaries of a linear fit ((Kloosterman et al., 2014); see **Methods**; decoded trajectory seen as a superimposed dotted white line in **Figure S1A,B** and examples in **Figure 3**). This captured probability, normalized by the total maximum theoretical probability that could be captured, is computed as the trajectory score, as previously described (Kloosterman et al., 2014; Drieu et al., 2018). We saw a substantial proportion of decodes with sequential, linear structure in both wake and REM sleep using this ensemble approach to delineate theta sequence events from non-sequence events (**Figure 3**).

**Figure 3.**
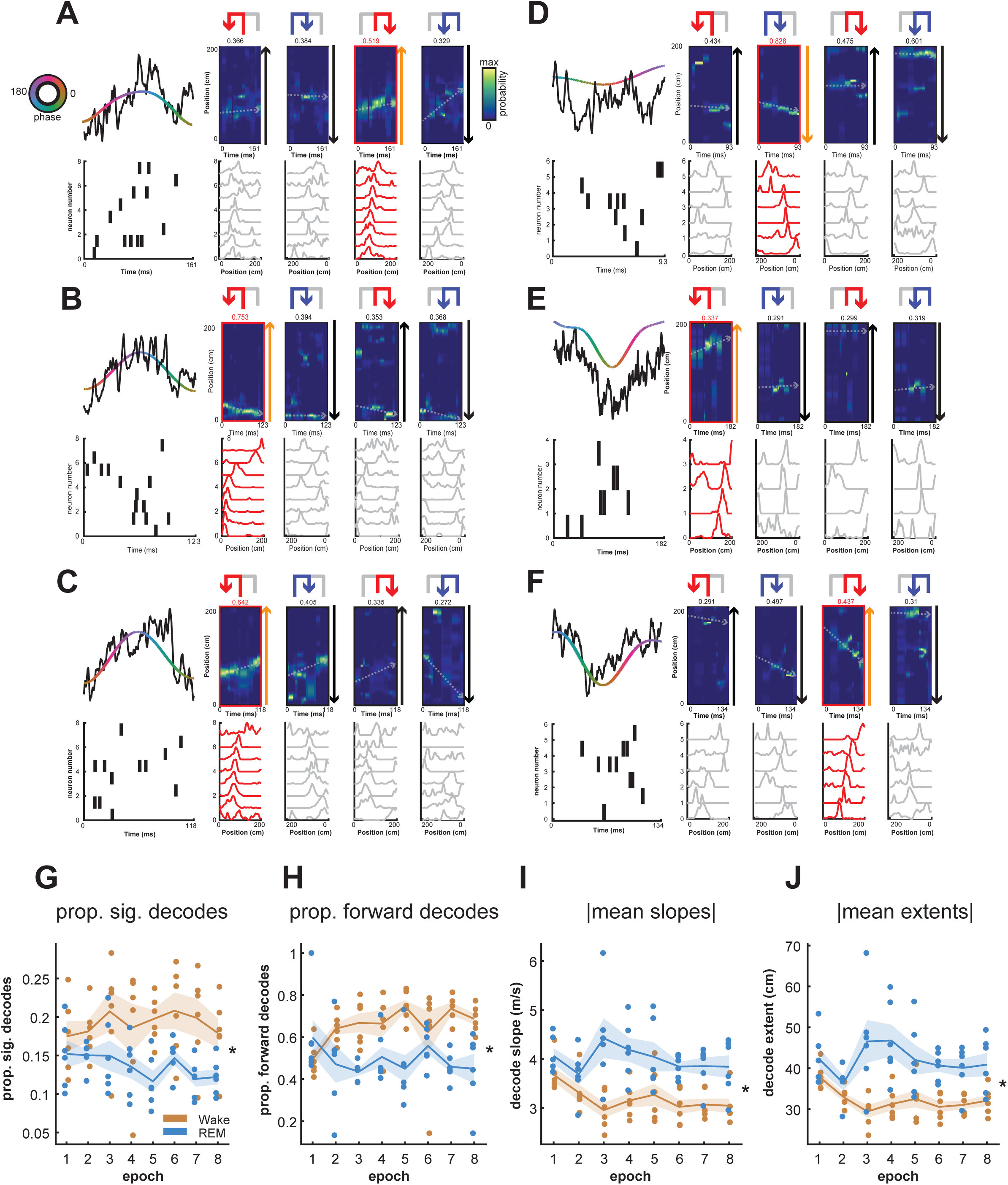
Theta sequence decoding and significance in wake and REM. (**A-F**) Example decoded theta sequences in wake (**A-C**) and REM (**D-F**). (**A**) Example decoded theta sequence in wake. This wake sequence significantly decodes an outbound right trajectory in the forward direction. **(Top Left)** The raw LFP trace of this theta cycle (black), and corresponding theta filtered LFP (rainbow indicating phase). **(Bottom Left)** Spikes and neurons in this theta sequence, sorted by peak firing of the chosen trajectory. **(Bottom Right)** Firing rate curves per trajectory type for the neurons at the bottom left, sorted and normalized according to peak firing of the chosen trajectory, highlighted in red. **(Top Right)** Decoded posteriors for each trajectory type, with warmer colors indicating higher spatial bin probability and corresponding putative theta sequence line fits overlaid. Arrows to the right indicate the animals’ direction of travel in this trajectory type, in order to differentiate forward and reverse decodes. Titles indicate normalized line fitting scores. Chosen trajectory and contributing firing rate curves highlighted in red, with direction of travel arrows in orange. (**B**) Same as previous. This wake sequence significantly decodes an outbound left trajectory in the reverse direction. (**C**) Same as previous. This wake sequence significantly decodes an outbound left trajectory in the forward direction. (**D**) Example decoded theta sequence in REM. This REM sequence significantly decodes an inbound left trajectory in the forward direction. (**E**) Same as previous. This REM sequence significantly decodes an outbound left trajectory in the forward direction. (**F**) Same as previous. This REM sequence significantly decodes an outbound right trajectory in the reverse direction. The inbound left trajectory was not chosen due to violation of the jump distance criterion. (**G**) Proportion of significant theta sequences over behavioral epochs for wake (orange) and REM sleep (blue). Wake theta sequences consistently had a higher proportion of significant decodes than REM sleep (p = 3.48e-07, Friedman Test). (**H**) Proportion of forward theta sequences over behavioral epochs for wake (orange) and REM sleep (blue). Wake theta sequences consistently had a higher proportion of forward decodes than REM sleep (p=1.49e-06, Friedman Test) (**I**) Mean absolute theta decode slopes over behavioral epochs. REM theta sequences consistently had a higher decode slope (p = 2.29e-10, Friedman Test). (**J**) Mean absolute theta decode extents over behavioral epochs. Similar to mean absolute decode slopes, REM theta sequences consistently had a higher decode extent (p = 2.29e-10, Friedman Test). Data in **(G)-(J)** are means (solid lines) ± SEM (shaded area). \**P* < 0.05; ns, not significant.

Previous literature has advocated for this ensemble approach to identify sequential reactivation events (Foster, 2017). We therefore used a rigorous combination of trajectory scores, shuffles, and other complementary measures to separate statistically significant decodes from candidate events and to reduce the risk of false positives, and applied the same method to both wake and REM states. Line fitting scores were overall higher in wake as compared to REM sleep (**Figure S1C**,wake and REM sleep distributions are different, p < 1e-99, rank-sum test), indicating the higher fidelity of decoded sequential positions and putative theta sequences in wake. Fitted trajectory slopes (**Figure S1D**) were significantly higher in REM decodes than in wake (wake and REM sleep distributions are different, p < 3.41e-59, rank-sum test), indicating higher virtual decode velocities, which were also mirrored in the absolute decode distances elapsed in these decodes (**Figure S1E**) and their differences (wake and REM sleep distributions are different, p < 1.46e-47, rank-sum test). Though recent work has provided a fascinating possible role for stationary representations of space in theta decodes (Muessig et al., 2019), we imposed virtual decode velocity cutoffs of greater than 1m/s and less than 10m/s to only analyze non-stationary decodes, or decodes with velocities approaching those reported of sharp-wave ripple replay to avoid potential, though unlikely false negatives.

Similar to reported methods from prior studies analyzing theta sequences (Zhang et al., 1998; Davidson et al., 2009; Karlsson and Frank, 2009; Gupta et al., 2012; Pfeiffer and Foster, 2013), we employed two shuffle procedures on decoded spatial posteriors, permuting both posterior time bin order (**Figure S1F**) or circularly permuting spatial bins within each time bin (**Figure S1G**) 1000 times, computing optimal line fitting trajectories for each permutation. The time bin shuffle or space shuffle scores were then computed as the percentile of the real. We imposed a 99^th^ percentile cutoff to ensure that putative theta sequences had scores distinguishable from those possible by chance alone (p < 0.01 criterion). Wake and REM sequences had small but significant differences in shuffle scores (**Figures S1F-G**; wake and REM sleep distributions are different, both p < 3.28 e-33, rank-sum test).

We also employed two metrics on decoded posteriors independent of trajectory score that impose both regularity and continuity of decoded sequential position, limiting both jump distance and intra-decode silence. Jump distance (Silva et al., 2015) imposes decode regularity on a putative decode by limiting the distance a maximal decoded position can travel over successive time bins within a posterior. Here, we limit the distance a putative decode can travel within one time bin to 30% of the track (**Figure S1H**). Continuity is enforced on a putative decode by limiting the total number of silent time bins within the probability containing portion of a decode posterior.

In cases where more than one trajectory template of a putative sequence passed these statistical criteria, we chose the trajectory template that had the highest real trajectory score, for both wake and REM sleep. Using this rigorous ensemble of statistical methods applied equivalently to both states, we investigated the properties of detected theta sequences in wake and REM sleep.

### REM theta sequences mirror waking theta sequences

With a principled way of delineating theta sequences from putative spatial decodes for both wake and REM, we examined the representation of trajectories, directionality, and other properties of these sequences. **Figure 3A-G** show six example decoded theta sequences, with **Figure 3A-C** showing wake decodes, and **Figure 3D-F** showing REM decodes. Additional examples are shown in **Figure S2**. Each example shows place cell activity (**lower left**) during a single cycle of theta oscillation (broadband/theta LFP in **upper left**) for a candidate event. Templates for all four trajectories are shown as linearized place fields sorted by peak locations on the detected theta sequence trajectory (**bottom right**), with the chosen/detected trajectory highlighted in red. Bayesian reconstructions of position from place cell firing in these cycles (**upper right**) are overlaid with trajectory fits (gray/red lines), with trajectory scores indicated on top. To the right of each reconstruction are arrows indicating the direction of animal movement along the track, allowing inspection of which direction the spatial sequences propagate. The entire dataset of putative and selected theta sequences for both wake and REM are shown using an unsupervised non-linear dimensionality reduction technique for visualization (UMAP, see *Interactive dataset visualization* in **Methods**), available at https://github.com/JadhavLab/ThetaSequencesInREM/.

We quantified the prevalence of theta sequences and their properties in wake and REM over the course of learning. The total proportion of significantly decoded theta sequences was significantly different for wake and REM sessions, with ∼14-17% for wake and ∼8-11% for REM (**Figure 3G**; There was a statistically significant difference between wake and REM theta proportions, χ^2^(1) = 25.96, p = 3.48e-07. Friedman Test.). A majority of these sequences were in the forward direction for wake, while the proportion of forward decodes in REM were balanced with those in the reverse direction, **Figure 3H** (proportion of forward theta sequences for wake vs. REM, χ^2^(1) = 22.62, p = 1.98e-6. Friedman Test). We also saw a consistent difference in mean absolute decode slope between wake and REM (**Figure 3I**; there was a statistically significant difference between wake and REM theta sequence slopes, χ^2^(1) = 40.21, p = 2.29e-10. Friedman Test.) and decode extent (**Figure 3J**; wake vs. REM theta sequence extents, χ^2^(1) = 40.21, p = 2.29e-10. Friedman Test).

### Theta sequences in wake and REM sleep propagate differentially in forward and reverse directions

In finding the optimal theta decode, we allowed any given trajectory or slope, and found that a large fraction of reverse theta decodes were chosen as being the most likely decoded event, especially in REM sleep which showed a balance between reverse and forward sequences, in contrast to predominantly forward sequences in wake (examples in **Figure 3, Figure S2**). Reverse theta sequences have been reported during waking behavior previously (Wikenheiser and Redish, 2013; Zheng et al., 2016; Drieu and Zugaro, 2019; Schmidt et al., 2019), but to our knowledge, their role in behavior is not clear, although a recent study has reported reverse and forward theta sequences within individual cycles representing past and future positions respectively during behavior (Wang et al., 2020).

In order to further quantify the robustness of this sequential decoding and to ascertain forward and reverse sequences, we employed an additional quantification metric known as the quadrant ratio, which uses the shape of the decoded posterior over time to capture the magnitude and direction of a decoded sequence. Here, the quadrant ratio imposes a second, more stringent criterion for each individual sequence, to further assess the statistical validity of decoded sequences with significant trajectory scores. To examine decoded waking sequences assessed by quadrant scores, average theta sequence decodes centered on animal position were generated over the course of behavior for both forward and reverse sequences (top and bottom in **Figure 4A**). These sequences show average structured sequential representations per epoch, with average LFP waveforms segmented by theta overlaid in white. Theta waveform asymmetry did not change significantly over learning for either wake or REM (**Figure S3A-D**).

**Figure 4.**
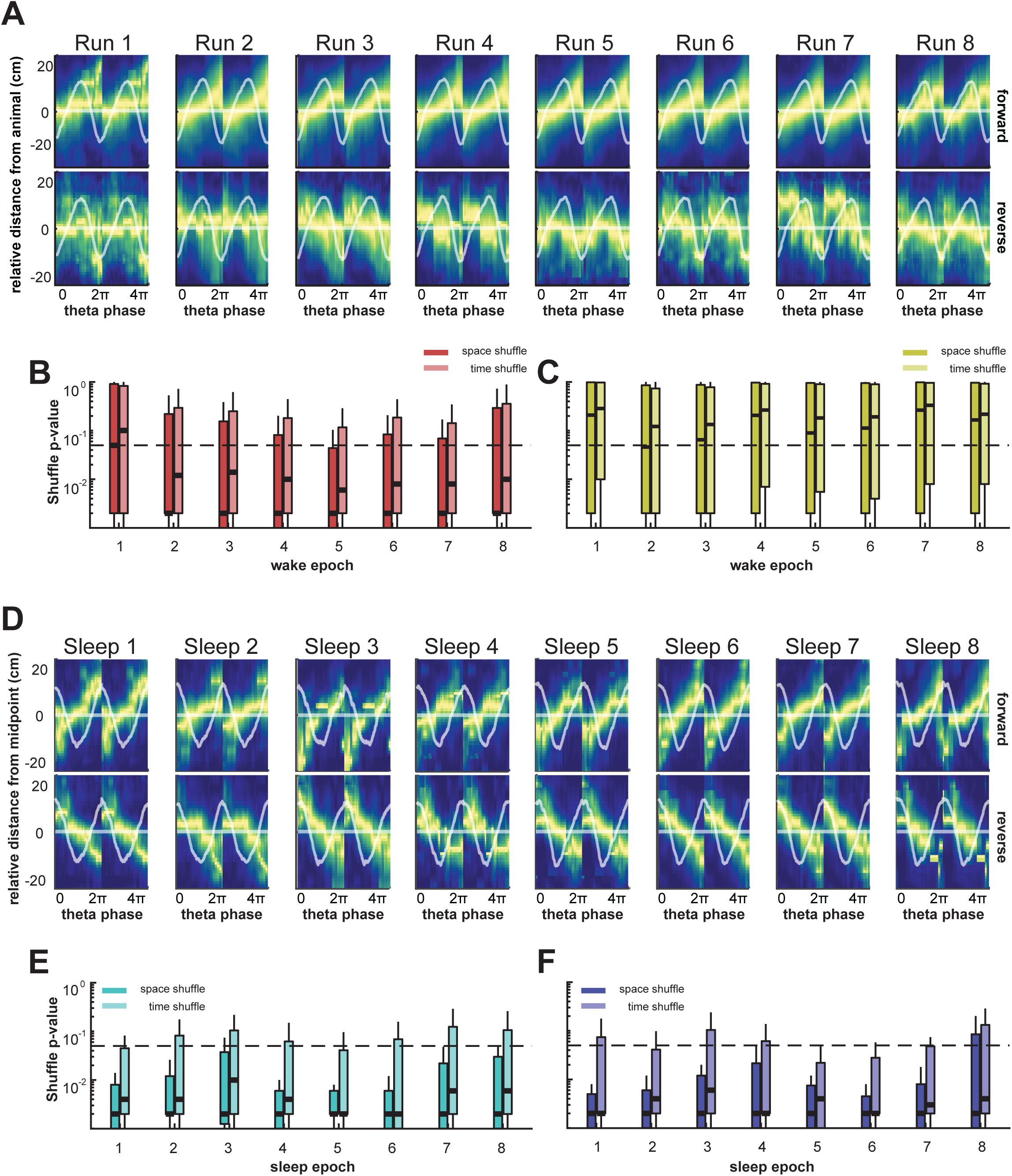
Theta sequence propagation in the forward and reverse direction. **(A)** Average theta sequences with respect to animal position in the forward **(top row)** and reverse direction **(bottom row)** during run sessions over behavior. Heat indicates maximum decoded probability over theta decode bins and relative distance from the animal position. Overlaid horizontal line indicates the decode midpoint, with overlaid mean LFP theta waveform and waveform SEM as shaded area. **(B)** Permutation test results for all forward theta sequences during wake, centered on animal position. Permutation tests were carried out using quadrant scores of individual decode posteriors versus the corresponding quadrant scores of randomly permuted decode posteriors, in order to create quadrant score null distributions. Lighter and darker boxplots indicate distributions of p-values using two different permutation strategies, shuffling decode posteriors in both space (circular permutation of space in each decode time bin) and time (permutation of decode posterior columns), respectively. Median p-values of all decodes over all epochs and tests are lower than 0.05 (thick dashed horizontal line). **(C)** Same as in **B**, for reverse (quadrant score less than 0) theta sequences during wake, centered on animal position. Median p-values are consistently higher than 0.05, indicating significant reverse decodes are rare relative to permuted decodes as measured by quadrant score. **(D)** Same as in **A** but with respect to decode midpoint in the forward and reverse direction during REM sleep sessions over behavior. **(E)** Same as in **B**, for forward theta sequences during REM sleep, centered on decode midpoint. Median p-values are consistently lower than 0.05, indicating significant forward decodes relative to permuted decodes as measured by quadrant score. **(F)** Same as in **C**, for reverse theta sequences during REM sleep, centered on decode midpoint. Median p-values are consistently lower than 0.05, indicating significant reverse decodes in REM sleep relative to permuted decodes as measured by quadrant score.

Each individual sequence centered on current animal position was assessed for quadrant score significance using spatial and time bin shuffles. **Figures 4B-C** show significance values from permutation tests comparing the probability of true quadrant scores relative to their shuffled counterparts (see **Methods**) for all individual wake theta sequences over learning in the forward (**Figure 4B**) and reverse (**Figure 4C**) directions, respectively. Shuffled distributions for each decode consist of random circular permutations of space in each decode time bin (space shuffle), and randomly permuting time bins (time shuffle). The permutation tests show that quadrant ratios develop rapidly for forward sequences in wake, especially in the second behavior epoch. Quadrant score permutation tests for large majority of forward theta sequences aligned by current animal position were clearly significant relative to shuffled distributions (**Figure 4B-C**; p = 0.05 is indicated by the dotted line, and p-values for median scores indicated by thick horizontal line overlaid on shuffle distributions. Note that the majority of p-value distributions lie below the p = 0.05 threshold for forward sequences from epoch 2 onward), indicating rapid formation of significant forward theta sequences early in behavior relative to shuffled controls. In contrast, only a small subset of individual reverse sequences in wake are significant for the quadrant score statistic (**Figure 4C**; with p = 0.05 indicated as a dotted line). Reverse theta sequences initially comprised a smaller proportion of wake theta sequences as assessed by the trajectory score metric (**Figure 3H**), and this quadrant ratio metric further suggests that significant reverse sequences are not very prevalent as compared to forward sequences during behavior.

We performed a similar quantification (**Figure 4D-F**) for REM sleep. Average theta sequences relative to current animal position and corresponding quadrant ratio scores are only valid for wake theta sequences. Since REM theta sequences were detected using waking trajectory templates and do not correspond to current animal position on the maze trajectories, this necessitated centering of decodes at the midpoint for quadrant score computation in REM (**Figure 4**). REM theta sequences were centered around decode midpoints, and again split into forward and reverse sequences. Quadrant score permutation tests were significant relative to both shuffle controls throughout learning for majority of REM sequences in either direction, forward and reverse. This confirms results from **Figure 3** that REM theta sequences develop rapidly and exhibit compressed sequential reactivation of waking spatial behavior in forward and reverse directions. Since centering by decode midpoint rather than animal position results in a more structured quadrant score by design, a stringent evaluation of the quadrant score statistic for REM sequences similar to wake sequences is not feasible. An equivalent comparison for wake theta sequences centered by decode midpoint is shown in **Figure S4A-C**.

### Theta sequence content and participation of phase reversing cell ensembles

We next wanted to determine whether and how the content of these theta sequences changes over time, and whether phase shifting cells (**Figure 2**) had any role in these sequences. Given the known difference in afferent connections between superficial and deep CA1 neurons, and their phase reversal in REM sleep (Mizuseki et al., 2011; Valero et al., 2015; Oliva et al., 2016b, a), we quantified the proportion of sequences with phase shifting neurons (termed “shifter sequences”) relative to those without phase shifting neurons (termed “non-shifter sequences”, sequences with a threshold of 25% or more shifting cells were considered to be shifting cell predominant; qualitatively similar results were seen with a threshold of 50%, see **Methods**).

First, we found that wake sequences overall have relatively even proportions of shifting and non-shifting sequences seen throughout behavior, whereas REM sequences comprise overall more non-shifting sequences (**Figure 5A;** there was a statistically significant difference between the proportion of wake and REM shifter and non-shifter sequences, χ^2^(1) = 8.16, p = 0.0043, Friedman Test). In order to summarize the content of these sequences and ascertain the contribution of the phase-shifting cell participation, we generated quiver plots showing the extent of theta sequence decodes (**Figure 5B-C**) in the forward direction for wake (**top**), and forward and reverse directions for REM sleep (**bottom**). Examples of quiver plots for illustrative individual sessions in early and late learning epochs are shown in **Figure 5B-C** respectively, and distributions of decoded positions attributed to shifter vs. non-shifter sequences are shown as histograms of spatial bin counts in the topmost row in **Figure 5B-C**. Ensembles which had more than 25% of phase shifting cells participating are in red (“shifter sequences”, see **Methods**; qualitatively similar results were obtained using a 50% threshold). For REM sequences (**Figure 5B-C, bottom**), note the lack of phase-shifting cell sequences (non-shifter sequences) in the early epoch, and the emergence of shifter sequences in the late epoch.

**Figure 5.**
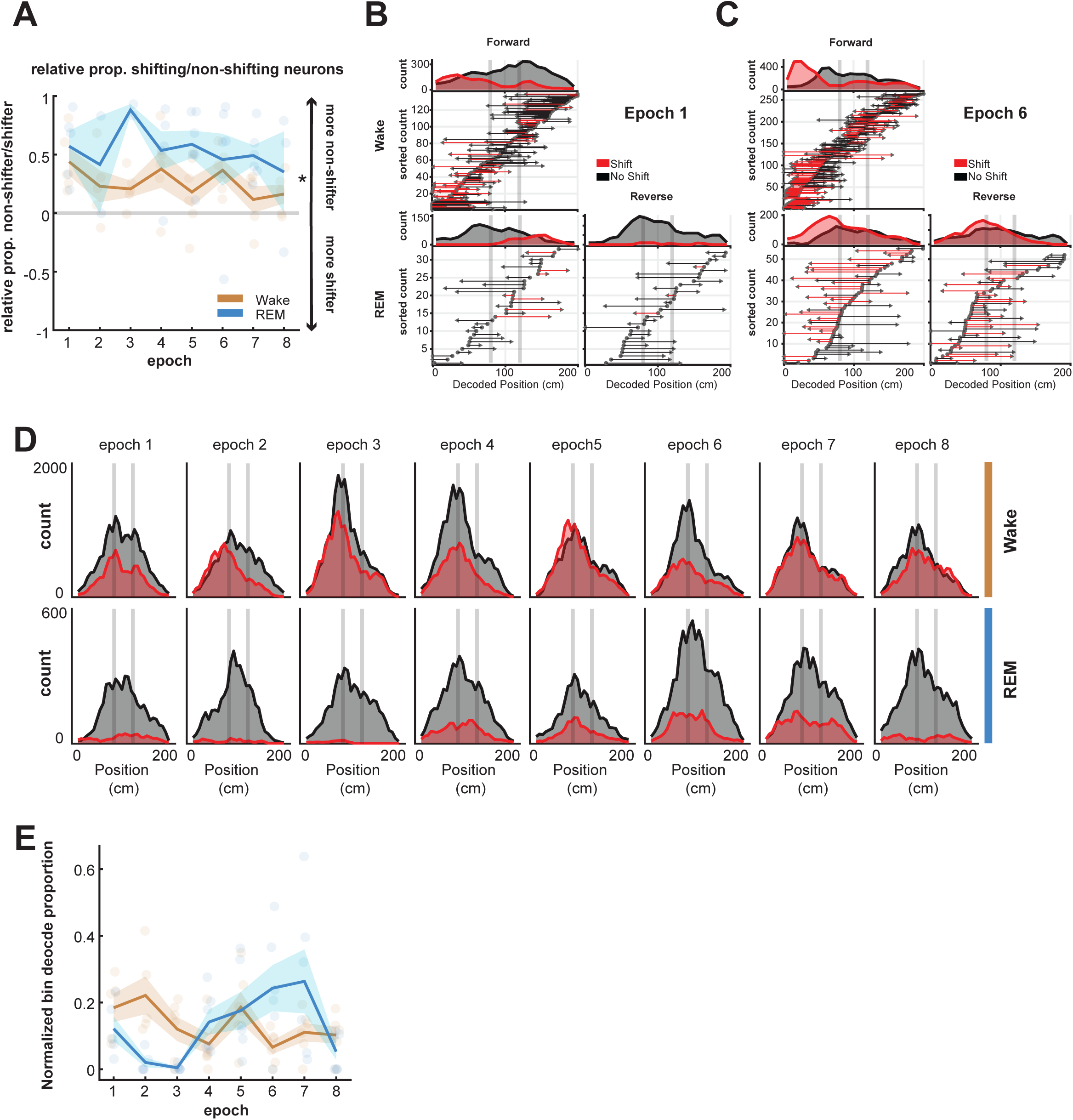
Theta sequence content and participation of phase-reversing cells. **(A)** Relative proportions of decodes comprising phase-shifting vs. non phase-shifting cells in wake (orange) and REM sleep (blue) over behavioral epochs. Decodes with 2 or more phase-shifting cells, which comprise of at least 25% of all participating place cells which shift their phase in REM sleep, are considered to be “shifter sequences” comprising shifting cells. Values greater than 0 indicate decodes with less phase shifting cells (or, more non-shifting cells); values near 0 indicate a balance between decodes with and without shifting neurons, and values below zero indicate decodes with more shifting neurons. Note the statistically significant difference between the proportion of wake and REM shifter and non-shifter sequences (p = 0.0043, Friedman Test). **(B)** Theta sequence trajectories for wake (Top) and REM (Bottom) sessions in the **(left)** forward and **(right)** reverse decode directions for an early learning session in one animal, sorted according to decode start position. Only forward sequences are shown in wake. Circles on the x-axis indicate decode starts, arrowheads on the x-axis indicate direction of decode ends. Lines in red indicate decodes with more than 25% reversing cells (termed “phase-reversing or shifter cell sequences”), while lines in black indicate less than 25% phase reversing cells. Top histograms indicate decode coverage of the track, split into phase shifting and non-phase shifting theta sequences. For REM sleep epoch, note dearth of phase reversing cell (red) sequences. **(C)** Theta sequence trajectories for wake (Top) and REM (Bottom) sessions in the **(left)** forward and **(right)** reverse decode directions for a late performance session. Only forward sequences are shown in wake. Note that REM sleep epoch has similar proportions of forward and reverse sequences and included many phase reversing cell (red) sequences. Note also increase in spatial coverage for phase reversing cell sequences in REM (red histogram in top row) for this late performance session as compared to early learning session in **A.** **(D)** Coverage histograms of all epochs, and behavioral states, with shifting sequence decodes in red and non-shifting sequence decodes in black. In REM, sequence decodes containing phase shifting neurons emerge over learning (note epochs 5-8), corresponding with asymptotic behavioral performance in run sessions. **(E)** Quantification of sequence decode coverage by shifter vs. non-shifter sequences over epochs for wake (**orange**) and REM (**blue**). Overall, there was a statistically significant difference in the normalized bin decode proportion for REM over epochs (blue, p=0.0016, Kruskal-Wallis test). Wake sequence decodes (**orange**) with phase shifting cells do not show this effect over epochs (p=0.0726, Kruskal-Wallis test).

We also examined spatial coverage of these decoded sequences across learning, quantified as histograms of spatial bin counts (summarized in **Figure 5D**). Decoded spatial coverage shows increase of track representation by shifter sequences in REM over learning (**Figure 5D,E**). We thus found that in REM, the contribution of sequences comprising phase-reversing cells to decoded locations on track, relative to sequences with predominantly non-phase reversing cells, increases over learning (**Figure 5E**; there was no statistically significant difference in the normalized bin decode proportion of shifter vs. non-shifter for wake sequences over epochs, χ^2^(7) = 12.98, p = 0.0726. Kruskal-Wallis test; there was a statistically significant difference in the normalized bin decode proportion of shifter vs. non-shifter for REM sequences over epochs, χ^2^(7) = 23.2, p = 0.0016. Kruskal-Wallis test).

## DISCUSSION

Our results establish the existence of theta sequences in REM sleep. We show that during spatial learning, REM theta sequences recapitulate time-compressed behavioral activity in the forward and reverse directions. REM theta sequences mirror the evolution and content of theta sequences in waking behavior at the trough of the theta cycle, with REM sleep exhibiting a balance of forward and reverse sequences in contrast to predominantly forward wake theta sequences. Finally, REM phase shifting and non-shifting cells, i.e. putative superficial and deep CA1 cells (Mizuseki et al., 2011; Poe, 2017), differentially participate in this evolution over behavior. Phase-shifting cells showed an increase in contribution to REM theta sequence representation after acquisition of learning. The existence of theta sequences in REM sleep, their development over learning of a memory-guided behavior, and the increased participation of a subset of phase reversing CA1 neurons in these sequential representations after learning, together point to REM theta-mediated sequences as another potential mechanism in hippocampally driven memory processes.

We find two notable differences in REM and wake theta sequences – balance of forward and reverse sequences, and differential participation of phase-shifting and non-phase shifting cells. The difference in the balance of forward and reverse theta sequences in waking and REM sleep can potentially be explained by the fact that waking circuits may be biased or shaped by incoming sensory input from entorhinal cortex and concurrent knowledge about the task structure from CA3. Forward theta sequences predominate in wake, and several previous studies have reported their role in deliberation, heading and goal representation (Johnson and Redish, 2007; Gupta et al., 2012; Cei et al., 2014; Wikenheiser and Redish, 2015a; Papale et al., 2016; Redish, 2016). Reverse theta sequences during wake have been reported (Wikenheiser and Redish, 2013; Zheng et al., 2016; Drieu and Zugaro, 2019; Schmidt et al., 2019), but to our knowledge, their role in behavior is not clear. A recent study used high-density hippocampal recordings to report the existence of reverse and forward theta sequences within individual cycles representing past and future positions respectively during behavior (Wang et al., 2020). This study also reported that REM phase-shifting cells are more likely to participate in sharp-wave ripple (SWR) associated reverse replay in sleep, suggesting a link to SWR-mediated consolidation.

Here, we observed a balance of forward and reverse theta sequences in REM that recapitulate recent experience in a time-compressed manner. The neuromodulatory landscape of REM shares an elevation of cholinergic tone with waking behavior, thought to be driven in part by the medial septum and other circuitry (Müller and Remy, 2017). A notable difference between these states are low levels of noradrenergic and serotonergic signaling, along with bursts of dopaminergic signaling in the hippocampus during REM sleep (Mizuseki and Miyawaki, 2017; Poe, 2017; Samanta et al., 2020). Interestingly, activity flow of the theta rhythm has been reported to reverse in REM via GABAergic mechanisms, propagating from the subiculum to CA3 exclusively during the neuromodulatory environment of REM sleep. Reverse theta sequences may thus be enabled by the unique neuromodulatory tone during REM sleep (Jackson et al., 2014; Genzel et al., 2015), and possibly play specific roles in memory.

The differential participation of phase-shifting cells in REM theta sequences over learning also suggests a specific role in memory processes. REM phase-shifting and non-phase shifting cells are known to be anatomically differentiated. The radial axis within CA1 is delineated in part by the differences in afferent connectivity, both external to the hippocampal circuit via MEC and LEC input to differing apical dendrites (Li et al., 2017; Masurkar et al., 2017), and within the hippocampal circuit, via CA2 and CA3 input to deep and superficial basal dendrites, respectively (Valero et al., 2015; Oliva et al., 2016b, a). This in turn informs their functional properties; place fields from superficial cells are less numerous and more stable, while deep cells have more place fields, which are flexible and encode task features, landmarks, and rewards (Mizuseki et al., 2011; Danielson et al., 2016; Geiller et al., 2017).

We observed differential prevalence of phase-shifting cells in REM vs. wake theta sequences, and increased participation of these phase-shifting cells in REM sequence representations after learning had occurred. This change in phase-shifting cell participation was selectively seen only in REM sequences and not wake sequences. Although there is a possibility that more shifting-cell theta sequences were simply detected later in learning based on the statistical methods employed, this is unlikely since the number of phase-shifting cells did not change across epochs, and the same statistical criteria were employed throughout learning for both wake add REM sequences. Rather, the emergent participation of phase-shifting cells in theta sequences suggests a potential role in memory processes, with a likely function being forgetting through depotentiation, as hypothesized previously (Poe, 2017).

Since firing of place cells at the theta trough is known to induce LTD at synapses (Huerta and Lisman, 1993, 1995; Hölscher et al., 1997; Orr et al., 2001; Griffin et al., 2004), it has been hypothesized that firing of phase-shifting cells at theta troughs in REM sleep can support depotentiation for circuit remodeling and forgetting (Poe, 2017). Such depotentiation may support new learning by weakening old associations and subsequently enabling new associations, or have a role in sparsening the contextual ensemble code for an environment as a means to generating a simplified code from an initial over-representation of environmental variables (Rasch and Born, 2013; Poe, 2017; Samanta et al., 2020). Our finding of REM sequences at theta troughs and the increased participation of phase-shifting cells in REM sequences after learning is suggestive of a possible mechanism for ensemble pattern sparsening through depotentiation of phase-shifting cells in REM. We speculate that if phase-shifting hippocampal neurons contribute to new and labile episodic representations, then the REM neuromodulatory environment can allow for de-potentiation and sparsening of newly formed synapses through theta sequences propagating in the forward and reverse direction. Labile representations mediated by REM phase-shifting hippocampal neurons can preferentially fire at the theta trough to eliminate any non-essential patterns of activity, leading to circuit remodeling and simplification of the contextual ensemble code. Theta sequences in REM can thus allow for exploration of novel recombination of experience with forward and reverse sequences, and simultaneous de-potentiation of selective sequences. Since phase-shifting cells preferentially participate in SWR reverse replay (Wang et al., 2020), the possibility of interactions between reverse theta sequences in REM and reverse SWR replay sequences in NREM sleep also cannot be ruled out. These hypotheses can be tested in future studies that examine the role of phase-shifting cells in learning multiple novel experiences, including reversal learning (Poe, 2017). Our findings thus support a potential role of REM sleep in memory formation, and establish the existence of bi-directional REM theta sequences as an integral component of the machinery involved in memory processes in sleep.

## ACKNOWLEDGEMENTS

This work was supported by NIH Grant R01 MH112661 to SPJ. Analyses were performed using Brandeis University’s High Performance Computing Cluster which is partially funded by DMR-MRSEC 1420382.

## AUTHOR CONTRIBUTIONS

M.C.Z. designed and performed all data analyses; J.D.S. performed experiments; S.P.J. conceived the study and supervised all aspects of the work; M.C.Z. and S.P.J. wrote the manuscript with input from all authors.

## METHODS

### Animals and experimental design

Six adult male Long Evans rats (450-550 g, 4-6 months, RRID: RGD_2308852) were used in this study. Data from four of these animals during behavior sessions has been reported previously (Shin et al., 2019). All procedures were conducted in accordance with the guidelines of the US National Institutes of Health and approved by the Institutional Animal Care and Use Committee at Brandeis University. Animals were housed individually and kept on a standard 12h/12h light-dark cycle with *ad libitum* food and water available. Daily handling and habituation took place prior to training and experimental protocols, with all protocols carried out during animals’ light cycles. As previously described (Jadhav et al., 2016; Tang et al., 2017; Shin et al., 2019), animals were food deprived to 85-90% of their stable *ad libitum* weight and trained to seek liquid food reward (evaporated milk) alternating between elevated linear track ends (80 cm, 7 cm wide track sections). Animals were also exposed and habituated to an enclosed and opaque elevated rest box during this training period over multiple days. After animals reached a criterion level during the linear track protocol (50 rewards in 15-20 minute linear track sessions), they were taken off food restriction and chronically implanted with a multi-tetrode drive (see *Surgical procedures, euthanasia, and histology*). Following recovery, animals were again food deprived to 85-90% of their *ad libitum* weight and re-habituated to the linear track task. Animals were then exposed to the full novel W-track continuous alternation behavioral task during the recording day (see *Behavioral task*), with electrodes positioned appropriately in the stratum pyramidale layer on the previous day.

### Behavioral task

Rats performed a continuous W-track spatial alternation task, as previously described (Jadhav et al., 2012; Jadhav et al., 2016; Tang et al., 2017; Maharjan et al., 2018; Shin et al., 2019). An experimental day consisted of multiple interleaved behavioral sessions (8 sessions) on the W-track and inactive periods in the elevated rest box, consisting of 15-20 min run sessions and 30-40 min rest sessions, respectively (**Figure 1A, B**). The W-track (∼80 x 80 cm) consists of elevated track sections (7 cm wide) with reward wells situated at the end of each of the three arms. The three arms were connected with two short sections (∼40 cm long) for a total of ∼200 cm for a behavioral trajectory from the center arm reward well, to the outer arm reward well. Calibrated evaporated milk rewards were delivered automatically via infrared detectors integrated in reward wells for correct trials. Rats were tasked with learning a continuous spatial alternation strategy (starting from the center arm), alternating visits to either side well (outbound component) and the center well (inbound component). Incorrect alternations (visiting the same side well in consecutive outbound components – outbound error), incorrect side-to-side well visits (without visiting the center arm – inbound error), or perseverations (repeated visits to the same well just visited) were not rewarded. Only correct trials were used for further analysis.

### Surgical procedures, euthanasia, and histology

Surgical implantation procedures were as previously described (Jadhav et al., 2012; Jadhav et al., 2016; Tang et al., 2017; Shin et al., 2019), and post-operative analgesia administered for several days post implantation, along with free food and water. Briefly, each rat was surgically implanted with a 3D printed microdrive containing 30 independently moveable tetrodes, with 15 tetrodes targeting right dorsal hippocampus (−3.6 mm AP and 2.2 mm ML) and 15 tetrodes targeting right PFC (+3.0 mm AP and 0.7mm ML; not considered here). Tetrodes were made by twisting and bundling 4 NiCr wires (diameter 13 μm; Sandvik Palm Coast, Palm Coast, FL), followed by gold electroplating to an impedance of 200-300 kOhm. Electrodes were gradually advanced for 2-3 weeks following surgery to desired depths, after recovery and concurrent with pre-training. The hippocampal formation and advancement into the pyramidal layer was identified by characteristic LFP patterns such as presence of sharp-wave ripples (SWRs), SWR polarity and theta modulation. At the end of the experiment, 24 hours prior to euthanasia, animals were anesthetized (1-2% isoflurane) and a current (30 µA) was passed through each tetrode to form lesions at their tips for localization. Animals were later euthanized (Beuthanasia 200 mg/kg) and perfused transcardially with 4% formaldehyde using approved procedures. Brains were then fixed in 4% formaldehyde and 30% sucrose, cut into 50-µm sections, stained with cresyl violet, and imaged for verification of lesions indicating tetrode localization.

### Electrophysiology and data acquisition

All tetrodes were referenced with respect to a cerebellar ground screw. For each animal, one tetrode in corpus callosum above the hippocampal layer served as a hippocampal reference electrode. All behavioral and electrophysiological data was acquired using a SpikeGadgets system (SpikeGadgets, San Francisco, CA). Digital electrophysiological data was acquired using 128-channel digitizing headstages, sampled at 30 kHz and saved to disk, with spike data bandpass filtered between 600 Hz and 6 kHz, and local field potential (LFP) bandpass filtered between 0.5 Hz to 400 Hz and down sampled to 1.5 kHz. Input and output triggers for behavioral and environmental data (eg. reward delivery) were recorded at 1-ms resolution and synchronized to electrophysiological data. Animal movement and behavior was recorded and tracked using an overhead color CCD camera (30 fps), with animal head position, speed, and orientation indicated by color LEDs affixed to the headstage apparatus and microdrive. Cameras were calibrated to provide a resolution of 0.1 cm/pixel, and spatial extent of LEDs permitted a tracking resolution of ∼2 cm.

### Unit identification and inclusion

Spike peaks in electrophysiological recordings were identified by a threshold crossing of 40 µV in the filtered spike band for hippocampus (CA1). Spikes were then manually sorted into putative units as previously described (Jadhav et al., 2012; Jadhav et al., 2016; Tang et al., 2017; Shin et al., 2019). Briefly, candidate unit spiking had clustering parameters extracted (spike width on each channel, spike amplitude, and principal components), and were clustered using a custom Matlab (MathWorks, Natick, MA; RRID: SCR_001622) cluster visualization program (MatClust). Clusters were judged based on waveform shape, isolation distance, and lack of ISI violation. Only well isolated and stable putative excitatory units were included, with putative interneurons identified and excluded based on average firing rate ≥ 15 Hz and spike width criteria, as previously described (Jadhav et al., 2016; Tang et al., 2017; Shin et al., 2019). Further, only neurons which fired at least 100 spikes in each session were included for further analysis.

### Spatial maps and linearization

Two-dimensional occupancy-normalized spatial firing maps were calculated for each unit when the animals’ speed was greater than 3 cm/s, with spikes binned in 1 cm square bins and smoothed with a 2D Gaussian (2σ, 6cm wide), excluding spiking during high ripple power (>3 SD ripple band power, see *LFP collection and high-theta segmentation*). The linearized spiking activity of each cell was then computed by first assigning the rat’s linear position along the 2D skeleton of the four possible linear behavioral trajectories (center arm reward well to outer arm reward well for outbound trajectories, and the converse for inbound trajectories; (Frank et al., 2000; Jadhav et al., 2016)). Spiking and animal occupancy closest to each linear 2 cm bin on these four trajectories was then used to calculate the smoothed, occupancy-normalized linear firing rate for correct trajectories. A peak firing rate of 3 Hz or greater was required for a cell to be considered a place cell in CA1 (Jadhav et al., 2016). These linearized trajectories with occupancy-normalized firing rates were used in all subsequent decoding analyses.

### High-theta segmentation and REM state identification

LFP was band pass filtered in the delta (1-4 Hz), theta (6-12 Hz), and ripple bands (150-250 Hz) using zero phase IIR Butterworth filters. We determined envelopes and phases by Hilbert transform, and took the ratio of the theta to delta envelopes at each time point for every hippocampal tetrode. High theta periods during behavior were detected using criteria for theta power, running speed, and exclusion of sharp-wave ripples (SWRs). Specifically, behavioral high theta windows were assigned as uninterrupted time periods at least one theta cycle long (∼100 ms) when the smoothed (1σ, 8s long) mean theta/delta ratio exceeded 2, no SWRs were detected with a 3 SD threshold in the absolute power of the ripple band, and animal speed was greater than 3 cm/s. REM state identification proceeded in the same way, with the exception that animal speed could not exceed a threshold of 3 cm/s. REM states were identified as previously described (Kay et al., 2016; Tang et al., 2017), with candidate sleep periods identified as times with speed less than a threshold of 3 cm/s preceded by extended immobility periods (speed < 3cm/s) for at least 30 sec. REM states were identified as periods within candidate sleep periods with the same conditions for high theta/delta power and low ripple power as behavioral theta periods. LFP and position data for sleep state were also visually inspected for accuracy. Peak theta frequencies per animal and session were determined by computing the power spectrum for theta filtered LFP and taking the maximum peak in the theta range.

### Phase locking, phase reversing, and preferred theta phase

Theta LFP phase for hippocampal unit spiking was relative to a hippocampal reference tetrode located in corpus callosum, as previously described (Jadhav et al., 2016). For a given behavioral session (sleep or wake in a given behavioral epoch), all neurons with >100 spikes in valid theta times were used for computing phase sensitivity metrics (described above as speed during correct trials >3 cm/s and ripple band power <3 SD). Phase locking was computed using a Rayleigh’s z test for non-uniformity on the theta phases of the chosen candidate spikes in a given behavioral session. Those neurons which had a Rayleigh’s z p value of < 0.05 were considered to be phase locked in a given behavioral session and used for determining preferred population theta phase preferences. Only neurons which had at least one session in REM and wake were used for segmentation of phase reversing neurons in REM, classified as neurons with preferred mean theta phases centered on 230° and within 90° (preferred mean theta phases ≥ 140° and ≤ 320°) similar to (Mizuseki et al., 2011).

### Theta cycles and decoding

Within high theta time windows detected as described above, the troughs (0°) (or peaks(180°), in the case of REM sleep decoding) of the hippocampal theta-filtered LFP were identified and used to segment valid theta cycles, discarding cycles where phase was ambiguous or reset, as previously described (Johnson and Redish; Gupta et al., 2012; Feng et al., 2015; Wu et al., 2017). Due to the properties of CA1 pyramidal cells reversing their preferred phases in sleep (Mizuseki et al., 2011), REM theta sequences were segmented using a peak-to-peak method. Only theta cycles with at least 3 simultaneously active template cells were analyzed, and theta cycle decoding of place was implemented as previously described (Johnson and Redish; Gupta et al., 2012; Wu et al., 2017). A Bayesian decoder was used to calculate the probability of the animal’s location given the firing rate templates of the neurons that fired and their spikes that occurred in each time window, where time windows were sliding 20ms temporal bins with 5ms of overlap. (Zhang et al., 1998; Davidson et al., 2009; Karlsson and Frank, 2009; Gupta et al., 2012; Pfeiffer and Foster, 2013). Briefly, the probability of the animal’s position (*pos*) across all total spatial bins (*S*) given a time window (*t*), containing Poisson spiking (*spikes*) of independent units is

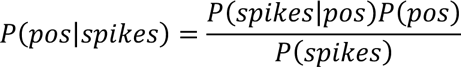

Normalizing over *P*(*spikes*) and using a uniform prior *P*(*pos*) to avoid spatial decoding bias, we get

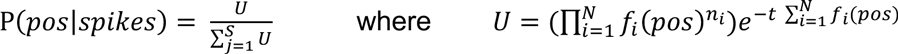

Where *f*_#_(*pos*) is the occupancy normalized 1D firing rate map for the *i*-th unit, *N* is the total number of active units, *n_i_* is the number of spikes fired by a particular *i*-th unit, and *t* is a time window (in this case, the entire theta cycle). This was computed using the firing rate template for the corresponding behavioral trajectories. The posterior then consists of a (*pos* x *t*) matrix, where *t* indicates the decoded time window, and *pos* represents the decoded space, with probabilities within each column *t* summing to 1. The “decoded” position per time bin was then the position with the maximum probability in that time bin.

### Theta Sequence Validation: Population Vector Overlap

In order to confidently differentiate decodes between different running directions in both wake and REM sleep, the similarity of the place-cell population between every behavioral trajectory in wake was computed using the population vector overlap (PVO) technique (Ravassard et al., 2013). The population vector (PV) was the group vector of all place cells in a certain linear position bin in a given epoch. The PVO was then defined as the vector dot product between the PVs across all linear positions in every epoch and every trajectory combination:

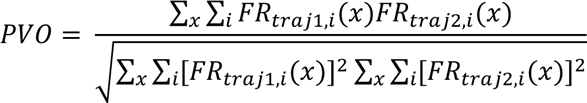

Where FR_traj1,i_(x) is the firing rate of the i-th place cell at the linear position x along the track on behavioral trajectory traj1, and FR_traj2,i_(x) is the firing rate of the i-th place cell at the linear position x along the on behavioral trajectory traj2. The PVO is a singular metric from 0 to 1, with 1 representing identical population place-field templates. To determine the significance of PVOs, we created a shuffled surrogate that permuted trial to trial assignments of spiking to behavioral trajectories for every epoch (behavior epochs 1 and 2 were combined due to low number of trials in epoch 1), repeating this procedure 1000 times and recomputing a null distribution to test our actual PVO values against. Data for sessions 1-2 for one animal was omitted from sequence analyses due to lack of sufficient number of trials.

### Theta Sequence Validation: Sequence Scores

For every putative theta sequence and every trajectory decoding template, the slope/line fitting procedure from (Davidson et al., 2009; Kloosterman et al., 2014; Feng et al., 2015) was used. Briefly, this decoding technique produces a line representing a putative theta sequence with a given slope (*V*) and intercept (ρ) maximizing the amount of probability captured within a 20-cm area around a line/theta sequence, fitted to the decoded posterior:

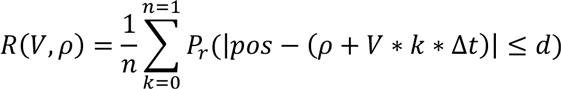

Maximizing the value *R*, where *Δt* is the moving window step (5 ms), d is 10cm (± 10 cm vicinity), theta time bins (*k*), and total time bin number (*n*). For time bins where a fitted line specifies a location beyond the track, the median probability of all likelihoods was chosen. Slope and intercept parameters were densely sampled to determine the values that maximized this sequence score.

### Theta Sequence Validation: Shuffles

Null distributions of theta sequence scores were determined as follows. Given a putative posterior containing a theta sequence, a null distribution was bootstrapped by randomly permuting this posterior 1000 times and computing the normalized sequence score as described above. Two permutation tests were carried out to construct two null distributions - randomly permuting the order of posterior columns and thus permuting time bins (henceforth known as time shuffle), and randomly circularly shifting columns and thus permuting space to which the posterior decoded (henceforth known as space shuffle).

### Theta Sequence Selection

After computing sequence scores for both null distribution shuffles and the true sequence score, the following was evaluated in order to select theta sequences from noisy or non-existent theta sequences. In practice, this selection required the following heuristics that imposed strict criteria to minimize false positives in theta sequence identification, at the risk of possible false negatives. A “true theta sequence” sequence score had to exceed the 99^th^ percentile of both space and time shuffled distributions. No more than 30% of the track could be traversed from one posterior time bin to another in the “decoded position” of the posterior, in order to enforce contiguity, used previously as a metric deemed the decode jump distance (Silva et al., 2015). No more than 3 bins (15 ms) of contiguous decoding ambiguity in the prior (time bins where no spikes occurred, or time bins where the posterior was uniform, and no maximally decoded spatial bin could be ascertained). This cutoff was used to enforce contiguity of decoding, rather than contiguity of decoded space. In addition, the virtual speed of the decode was restricted to the range of 1 m/s to 10 m/s, to avoid any possibility of stationary decodes, which have been described previously (Farooq and Dragoi, 2019; Muessig et al., 2019) but were beyond the scope of this analysis, as well as decodes that traversed the track faster than a single theta cycle, which were deemed to be artifactual or ripple-like given previous literature. In addition, only the tail ends of all sequence scores were considered, discarding any sequences whose true sequence scores were below the median of the population. With these cutoff criteria, what remains is determining which of the four decoding templates (OL, IL, OR, IR) is the correct decoding template. In practice, this could be determined by using the true trajectory the animal was currently on for awake theta sequences, but not during REM sequences. To ameliorate this, after these thresholding procedures, the decoding template that had the maximum true sequence score was chosen, similarly for both awake and REM theta sequences.

### Average theta sequences

Average theta sequences were constructed as follows. Each posterior of a selected theta sequence during awake behavior was shifted such that the position of the rat was at 0 cm, with theta sequence decodes for both decoding directions merged. The first 30 cm behind and ahead of an animal at a given theta sequence decode were then shown. For REM theta sequences, a similar procedure was carried out with some exceptions. Sleep decodes were aligned such that the theta cycle time midpoint was considered, and the decoded position at that timepoint of the sequence posterior was used to center the decode to 0 cm. Due to the nature of the slope-fitting maximization parameters, negative slopes were not restricted, that is, we allowed for theta sequences with negative slopes to be found in this maximization procedure. In practice, this means that we allowed for the possibility of a theta sequence to occur against the animal’s direction of travel for a given trajectory, i.e. reverse theta sequences.

### Average theta sequences-quadrant scores and validation

To quantify the structure of average theta sequences over time periods, we computed the quadrant ratio for each given posterior (Farooq and Dragoi, 2019). Briefly, the quadrant ratio gives a value from −1 to 1 indicating how much probability within the decoded posterior exhibits a canonical theta sequence, where the posterior is split into four quadrants, centered on the midpoint of the theta sequence on the x axis and centered along a relative position on the y axis. Quadrants II and III are located early in the theta cycle, while quadrants I and IV are located late in the theta cycle; while quadrants I and II are oriented in front of a spatial midpoint, and quadrants III and IV are located behind a spatial midpoint.

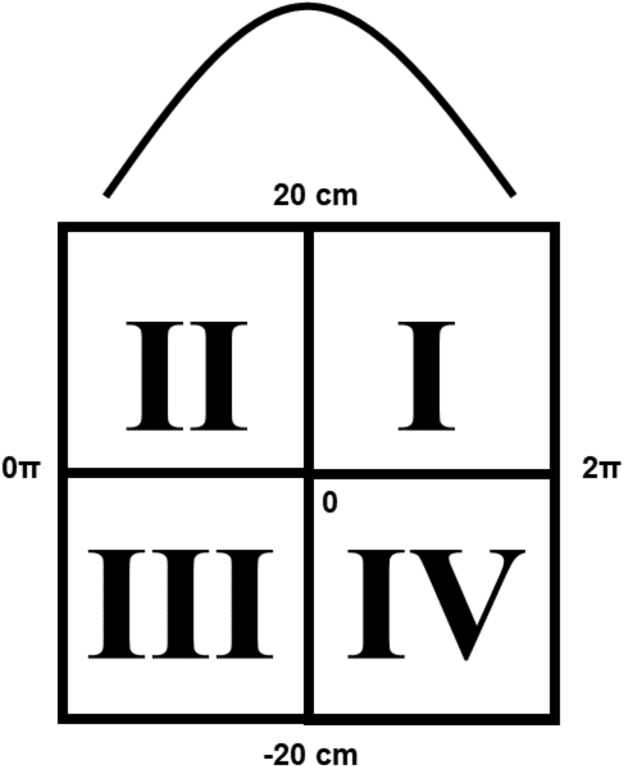

A canonical theta decode posterior exhibits the following structure, where a decode sweeps from behind the animal spatially early in a theta sequence (Quadrant III/QIII) to spatially in front of the animal late in a theta sequence (Quadrant I/QI) over time. Summing the total probability of the posterior in a given window that falls into QIII/QI, subtracting QII/QIV and normalizing by total probability in this window would lead to quadrant score values closer to 1, indicating a stronger likelihood of a structured forward theta sequence. A reversed theta sequence would then be ahead of the animal’s current position early in a theta sequence (Quadrant II/QII) and behind the animal’s current position late in a theta sequence (Quadrant IV/QIV), where values closer to −1 would indicate a strong likelihood of a structured backward theta sequence.

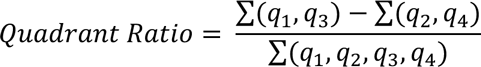

Each theta decode posterior at least 20cm from either end of the track was centered either according to the animal’s current spatial position (in wake), or midpoint of the putative theta decode (in REM), and then truncated to a local spatial area ± 20cm from this spatial midpoint for further quadrant score computation. Similar to validation of theta sequence decodes, the following procedure was carried out to create null distributions to test for significance. Each theta sequence posterior used for quadrant ratio quantification was randomly permuted 500 times, again permuting column order (time shuffle, permuting theta time bins) or circularly shifted along columns (space shuffle, permuting the spatial structure of the posterior per time bin), and quadrant ratios were recomputed. The real quadrant score for a given theta decode posterior was then compared to its corresponding null distribution in either shuffle type, and its percentile plotted.

### Quiver Plots, Cumulative Distributions, and Coverage Ratios

Since the line fitting procedure for quantification of theta sequences returns an assumed contiguous decode, we can also visualize the starts and ends of theta sequences, their spatial extents, and the distributions of decodes along the length of the tracks over time. To do this, we constructed quiver plots, as in (Wu et al., 2017), with the following properties. For a given theta sequence, an arrow is plotted where the *x*-axis indicates decoded positions-circles at the ends of arrows represent theta sequence decode start positions, and triangle ends represent theta sequence decode end positions. The *y*-axis position of the arrow delineates the ordinal value of the theta sequence, ordered according to decode start location on the track. Track arm extents are skeletonized with vertical lines indicating maze corners. To summarize the spatial extent of theta sequences, sequence extents are binned in 1-cm bins, then plotted as histograms of total decode extents, showing total spatial coverage of decodes, as well as any biases or over representation of space. Decodes with 2 or more phase-shifting cells, which comprise of at least 25% of all participating place cells which shift their phase in REM sleep, are considered to be sequences which had an ensemble of shifting cells. Qualitatively similar results were obtained using a 50% threshold.

To quantify the spatial coverage decodes in REM and wake by phase-shifting ensembles and non-shifting ensembles, we computed the coverage ratio between the two:

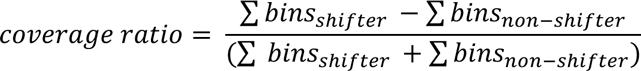

For each behavioral epoch, we subtracted the summed coverage in non-shifting ensembles from the coverage in shifting ensembles, normalized by the total number of covered bins in both ensembles, which resulted in a number varying from −1 (track over-representation by non-shifting ensembles) to 1 (higher track representation by shifting ensembles).

### Data Visualization

Colormaps used throughout were modified from the matplotlib package and adjusted with vscim, with the goal of being perceptually uniform (Thyng et al., 2016). Figure layout was generated in part using a ggplot2-like package created for MATLAB, Gramm (Morel, 2018).

### Data Availability and Interactive dataset visualization

The entire dataset of putative theta sequences for wake and REM is available for visualization at www.github.com/JadhavLab/ThetaSequencesInREM/. To visualize this dataset of putative theta sequences, we used an unsupervised non-linear dimensionality reduction technique known as Uniform Manifold Approximation and Projection (UMAP) (McInnes et al., 2018). Briefly, UMAP is a manifold learning technique that finds a lower dimensional embedding of higher dimensional data, much like t-SNE, though UMAP preserves both global and local structure (see https://umap-learn.readthedocs.io/en/latest/ for details). Each point indicates a single putative theta sequence prior to our selection criteria, color coded according to line-fitting score (warmer colors indicating lower line-fitting score, while cooler colors indicating higher line-fitting score, see colorbar for values). Each point, when hovered over, shows a decoded sequence, with x axis indicating time, y axis indicating track space, and color indicating probability of decode in that time x space bin (saturated for visualization). Drop down menus toggle between line-fitting score and selected/non-selected sequence color maps (selected decodes in dark blue, and non-selected decodes in yellow), respectively. An optional alpha/opacity dropdown can be used to indicate/highlight selected/non-selected decodes (to highlight selected decodes, choose the options, Values = selected; Alpha = selected_alpha).

## SUPPLEMENTARY FIGURE LEGENDS

**Figure S1.**
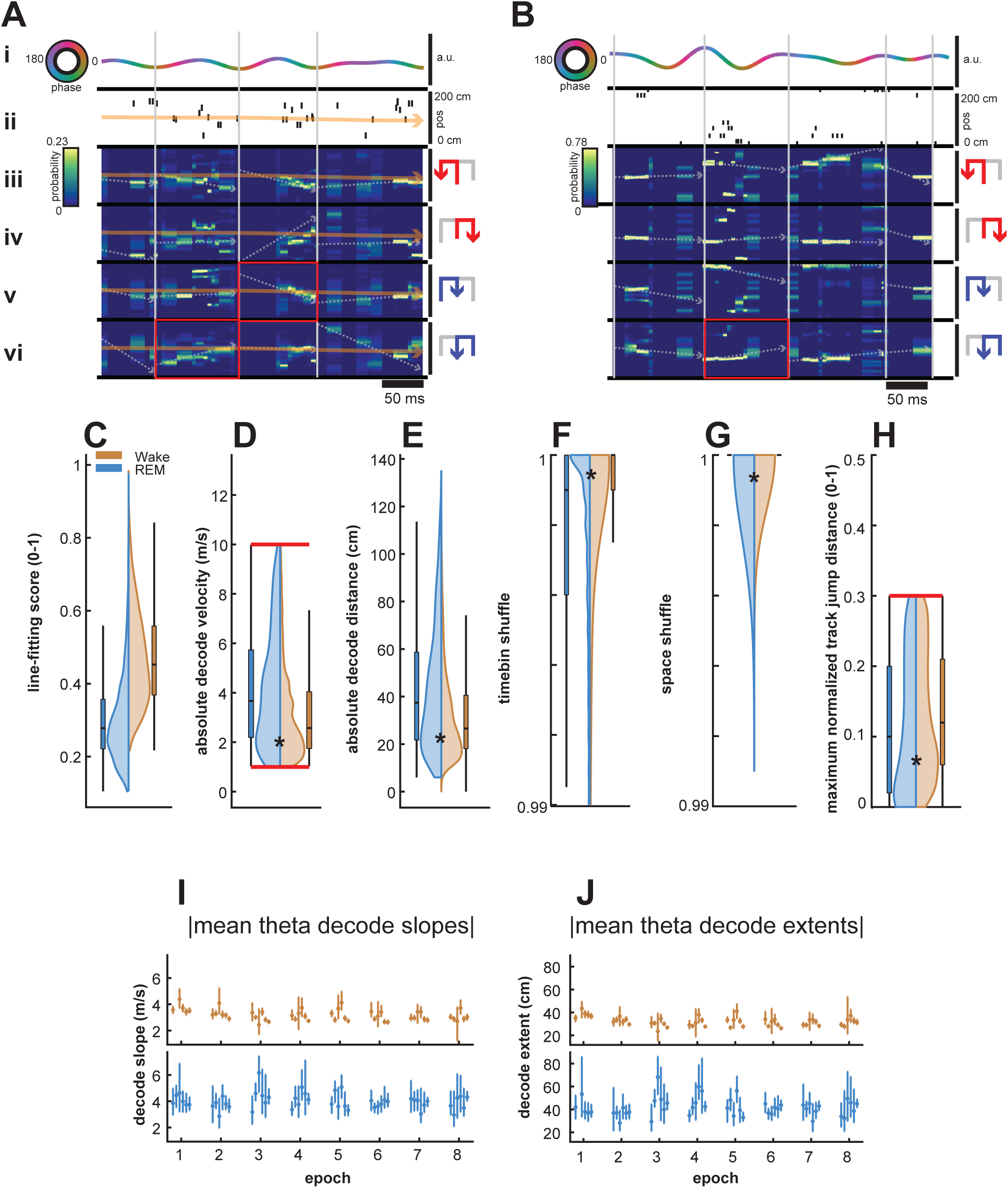
Theta sequences and their properties, Related to Figure 3. **(A)** 0.5 seconds of theta decodes during waking behavior. **(i)** Theta filtered LFP, scaled to maximum of the entire session, theta phase in (rainbow). Note parsing of cycles from trough to trough during wake**. (ii)** Spikes from all neurons, sorted according to peak field position along the track (y-axis, 0-200cm). Animal’s current position is shown by the orange line. **(iii)**-**(vi)** theta cycle decodes from trajectory templates ordered vertically as OL, IL, OR, IR respectively. Heatmap indicates probability, with the x-axis indicating decode time-bin, and the y-axis indicating decoded position (0-200cm), with the same position scale as **(ii)**. Line fits of putative theta sequences shown with dotted arrowhead lines (significantly decoded theta sequence shown as solid red line with arrowhead). Animal’s current position in orange, as in **ii**. **(B)** Same as **A**, 0.5 seconds of theta decodes during REM sleep. Note parsing of theta cycles from peak to peak **(i)**, and the lack of the animal’s current position **(ii)**. **(C)** Normalized line fitting scores, calculated as the summed captured probability of the putative theta sequence in each decode time bin divided by the summed theoretical maximal captured probability in each time bin (wake and REM sleep distributions are different, p <1e-99, rank-sum test). **(D)** Absolute virtual decode velocities of putative theta sequences. Only theta sequences with virtual velocities greater than 1 m/s or less than 10 m/s were included for further analysis (wake and REM sleep distributions are different, p <3.41e-59, rank-sum test). **(E)** Absolute decoding distance of all putative theta sequences in wake (orange) and REM sleep (blue), calculated as the absolute distance from the first decoded spatial bin to the final decoded spatial bin (wake and REM sleep distributions are different, p <1.46e-47, rank-sum test). **(F)** Time bin shuffle percentile. For a single given posterior representing a putative theta sequence, time bins (columns) were permuted 1000 times at random, and normalized line fitting scores were calculated for each of these shuffled posteriors. The time bin shuffle percentile is the percentile of the true normalized line fitting score in comparison to the distribution of the shuffled line fitting scores. Only theta sequences which exceeded the 99^th^ percentile were included for further analysis (wake and REM sleep distributions are different, p <7.96 e-54, rank-sum test). **(G)** Space shuffle percentile. For a single given posterior representing a putative theta sequence, posterior columns (representing decoded space) were circularly permuted 1000 times at random, and normalized line fitting scores were calculated for each of these shuffled posteriors. The space shuffle percentile is the percentile of the true normalized line fitting score in comparison to the distribution of the shuffled line fitting scores. Only theta sequences which exceeded the 99^th^ percentile were included for further analysis (wake and REM sleep distributions are different, p <3.28 e-33, rank-sum test). **(H)** Maximum normalized track jump distance, computed as the maximum track distance a putative decode travelled from one time bin to another. Only theta sequences with jump distances less than 30% of the track were included for further analysis (wake and REM sleep distributions are different, p <8.32e-20, rank-sum test). **(I)** Mean absolute theta decode slopes over behavioral epochs per individual animal. **(J)** Mean absolute theta decode extents over behavioral epochs per individual animal. Data in **(I)-(J)** are means (points) ± 95% CI (lines).

**Figure S2.**
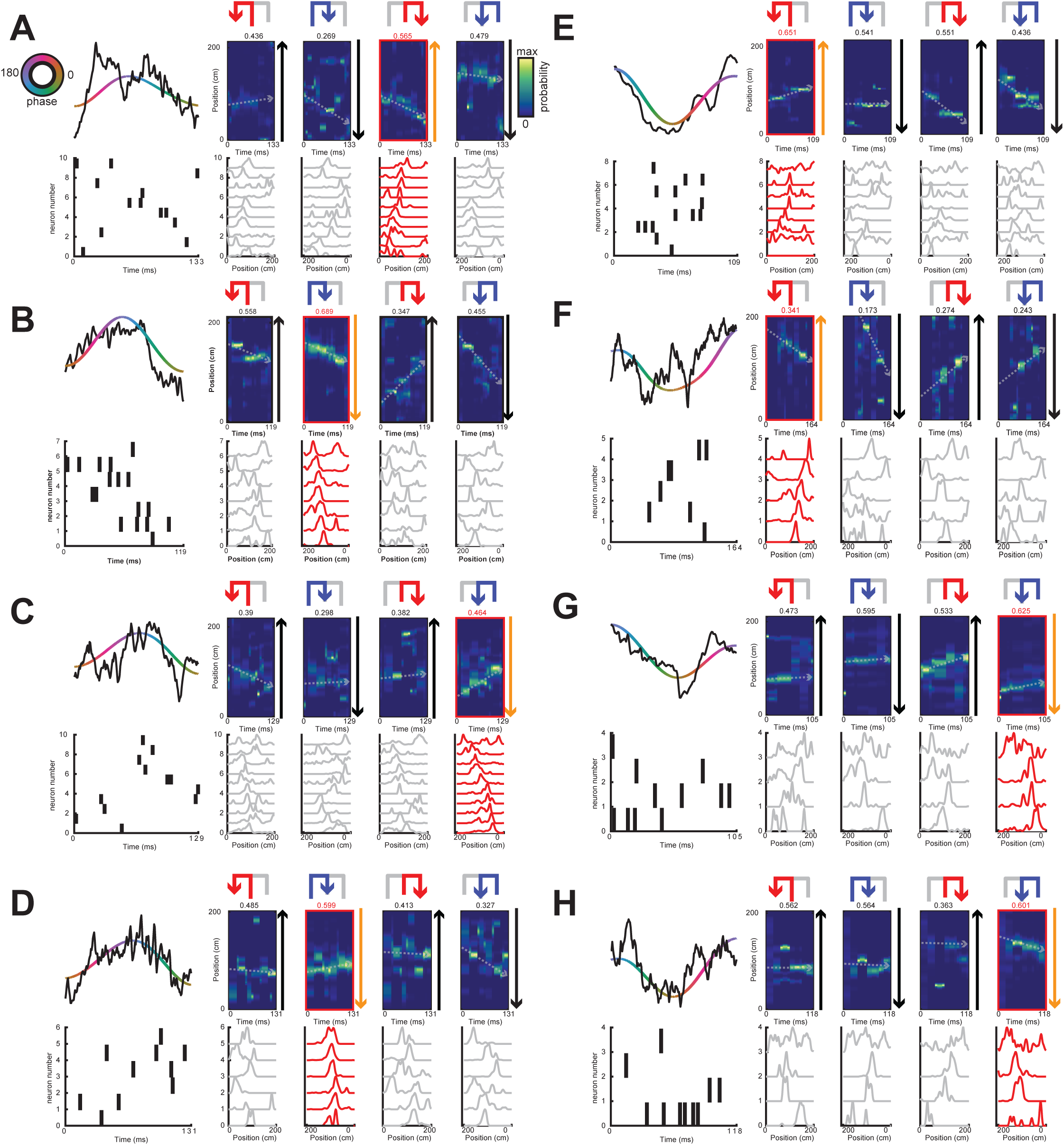
Additional theta sequence examples; Related to Figure 3 Same as Figure 3. **(A-E)** Examples of wake theta sequences **(F-J)** Examples of REM theta sequences

**Figure S3.**
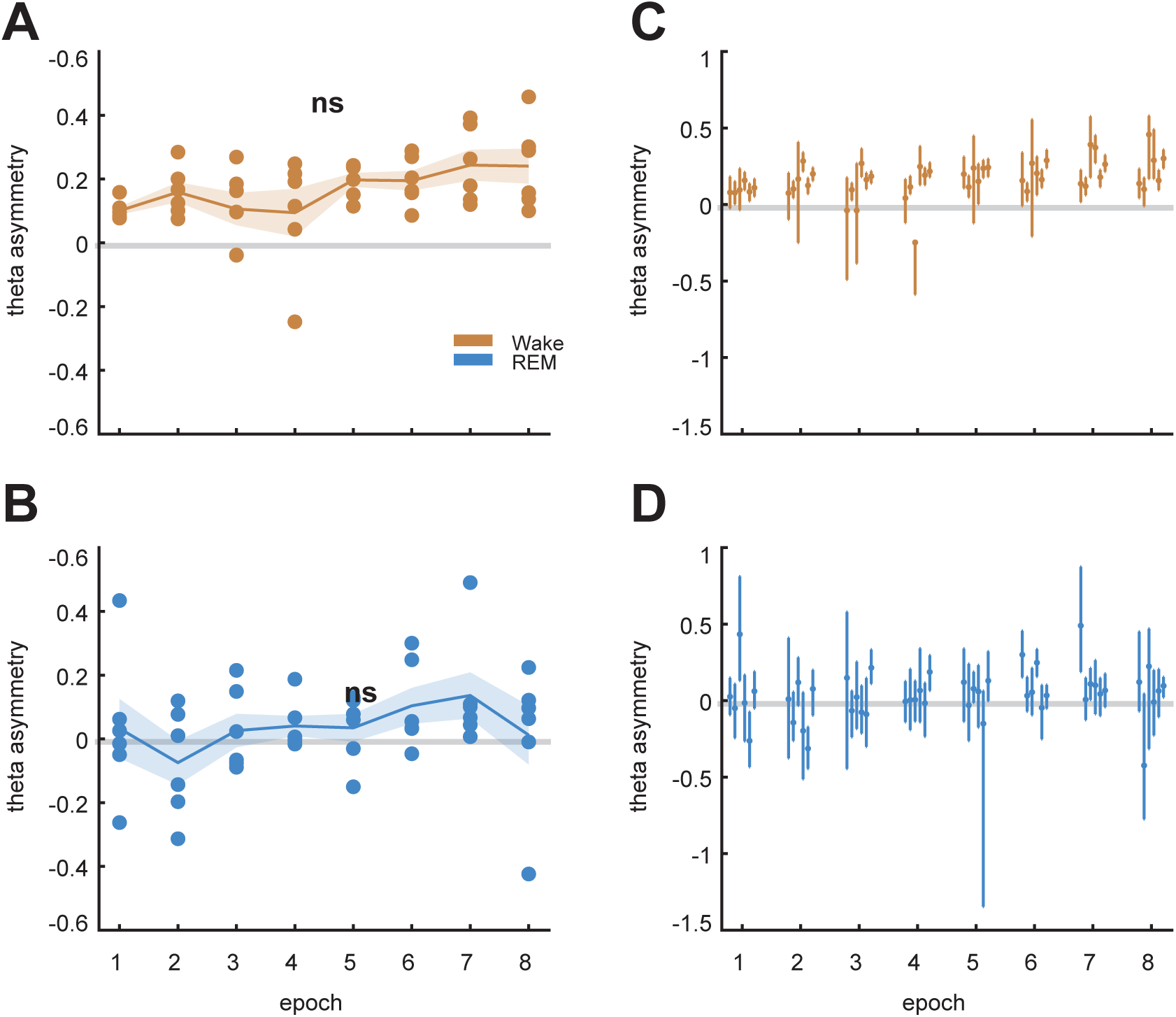
Theta asymmetry; Related to Figure 4. **(A)** Mean theta waveform asymmetries in wake are not significantly different over epochs (p=0.1054, Kruskal-Wallis test), with theta asymmetries in **(C)** shown as mean asymmetry score and CI per animal. **(B)** Mean theta waveform asymmetries in REM sleep are not significantly different over epochs (p=0.609, Kruskal-Wallis test), with **(D)** same as **(C)**. Data in **(A)-(B)** are means (solid lines) ± SEM (shaded area). \**P* < 0.05; ns, not significant. Data in **(C)-(D)** are means (points) ± 95% CI (lines).

**Figure S4.**
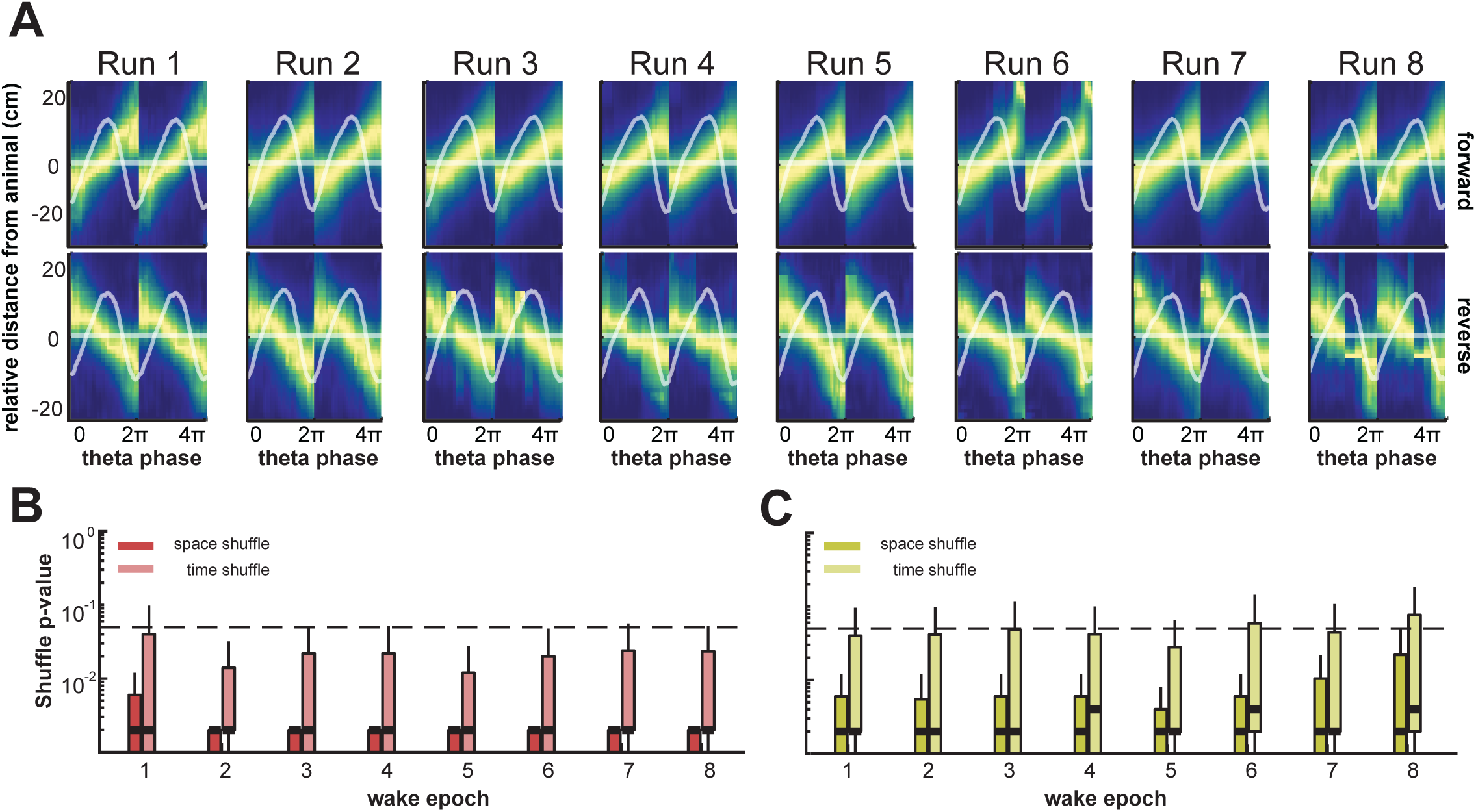
Wake quadrant score quantification according to midpoint alignment; Related to Figure 4. **(A)** Average theta sequences with respect to decode midpoint in the forward **(top row)** and reverse direction **(bottom row)** during run sessions over behavior. Heat indicates maximum decode probability over theta decode bins and relative distance from the animal. Overlaid horizontal line indicates the animal position, with mean LFP waveform and waveform SEM as shaded area. **(B)** Permutation test results, for forward theta sequences centered on decode midpoint during wake. Median p-values are consistently lower than 0.05, indicating significant forward decodes relative to permuted decodes as measured by quadrant score. **(C)** Same as in **B**, for reverse theta sequences centered on decode midpoint during wake. Median p-values are consistently lower than 0.05 indicating significant reverse decodes relative to permuted decodes as measured by quadrant scores.

